# Molecular Epidemiology of multidrug-resistant *Klebsiella pneumoniae, Enterobacter cloacae,* and *Escherichia coli* outbreak among neonates in Tembisa Hospital, South Africa

**DOI:** 10.1101/2023.12.15.571515

**Authors:** John Osei Sekyere, Masego Mmatli, Anel Bosch, Ramathetje Virginia Ntsoane, Harishia Naidoo, Sinenhlanhla Doyisa, Nontuthuko E. Maningi, Nontombi Marylucy Mbelle, Mohamed Said

## Abstract

**Background:** An outbreak of multidrug-resistant *Klebsiella pneumoniae, Escherichia coli, and Enterobacter cloacae* infections in a neonatal ward within a tertiary hospital in South Africa resulted in the mortality of 10 patients within six months. In this work, the genomic epidemiology of and the molecular factors mediating this outbreak were investigated.

**Methods:** Bacterial cultures obtained from clinical samples collected from the infected neonates underwent phenotypic and molecular analyses to determine their species, sensitivity to antibiotics, production of carbapenemases, complete resistance genes profile, clonality, epidemiology, and evolutionary relationships. Mobile genetic elements flanking the resistance genes and facilitating their spread were also characterized.

**Results:** The outbreak was centered in two major wards and affected mainly neonates between September 2019 and March 2020. Most isolates (n = 27 isolates) were *K. pneumoniae* while both *E. coli* and *E. cloacae* had three isolates each. Notably, 33/34 isolates were multidrug resistant (MDR), with 30 being resistant to at least four drug classes. All the isolates were carbapenemase-positive, but four *bla*_OXA-48_ isolates were susceptible to carbapenems. *Bla*_NDM-1_ (n = 13) and *bla*_OXA-48/181_ (n = 15) were respectively found on *IS*91 and *IS*6-like *IS*26 composite transposons in the isolates alongside several other resistance genes. The repertoire of resistance and virulence genes, insertion sequences, and plasmid replicon types in the strains explains their virulence, resistance, and quick dissemination among the neonates.

**Conclusions:** The outbreak of fatal MDR infections in the neonatal wards were mediated by clonal (vertical) and horizontal (plasmid-mediated) spread of resistant and virulent strains (and genes) that have been also circulating locally and globally.

## Introduction

*Escherichia coli, Enterobacter cloacae* complex, and *Klebsiella pneumoniae* are three of the most challenging Gram-negative bacterial pathogens implicated in numerous nosocomial infections and outbreaks ^1–3^. They are mostly associated with multidrug resistance in urinary tract infections, bacteraemia, meningitis, and pneumonia ^2,4,5^. Among infants, these two pathogens are mainly associated with blood-stream infections (sepsis) and pneumonia. Outbreaks among neonatal units and these bacteria are usually antibiotic resistant ^3^.

*E. coli, E. cloacae,* and *K. pneumoniae* have been associated with resistance to carbapenems, colistin, tigecycline, other β-lactams, fluoroquinolones, aminoglycosides, tetracyclines, fosfomycin, and all other antibiotic classes ^2,5–7^. It is therefore not uncommon for these species to harbour several resistance determinants and express multidrug resistance phenotypes. Furthermore, highly virulent strains, including hypervirulent *K. pneumoniae*, have emerged and several reports have shown the presence of multidrug resistance and hypervirulence in nosocomial and community *K. pneumoniae* strains ^8,9^.

From September 2019, physicians at Tembisa hospital, South Africa, began to see a spike in neonatal infections. The paediatrician notified the Department of Medical Microbiology at the University of Pretoria and requested an investigation into the cause of the spike in infection among neonates. Unfortunately, while preparations were ongoing to initiate the investigations, an outbreak occurred between November 2019 and January 2020^10^. Twenty infants were infected with carbapenem-resistant *K. pneumoniae* and *E. coli* during this outbreak and 10 demised. The infections however continued until March 2020^11^.

Clinical samples from the hospital’s neonatal unit that had been sent for diagnosis at the National Health Laboratory Services (NHLS), Tshwane Academic Division/Department of Medical Microbiology, University of Pretoria, between September 2019 and March 2020, were therefore collected and analysed. A series of phenotypic and molecular analyses of the samples were undertaken to delineate the molecular epidemiology and resistance mechanisms of the strains involved in the outbreak. Using whole-genome sequencing and phylogenomics, the evolutionary relationship between the isolates and other regional and global strains as well as the genetic context of their resistance determinants is characterised herein.

## Methods

### Study setting and samples

An uptick in carbapenemase-mediated infection outbreak was observed at the Tembisa hospital in September 2019. We thus followed up on this to trace and reign in the infection from further spread. Forty-five neonatal demographic data and 48 clinical samples from the Tembisa hospital were sent to the National Health Laboratory Services (NHLS), Tshwane Academic Division/Department of Medical Microbiology, University of Pretoria, between September 2019 and March 2020. The clinical and demographic data were curated into Microsoft Excel for downstream statistical analysis. The host age, disease, and sample sources were included in the clinical data. Descriptive statistics were used to analyse the demographic data.

### Isolate identification and resistance screening

The samples (48 stored isolates) were retrieved from the -70°C freezer, thawed and plated out on blood agar plates (BAP) (Diagnostic Media Products, South Africa) for further testing. However, only 40 isolates could be revived. Hence, all subsequent phenotypic and PCR tests were done on 40 isolates. The species and antibiotic resistance profiles of the forty isolates were identified using Vitek 2 (Biomerieux, Johannesburg, South Africa). Furthermore, the carbapenem minimum inhibitory concentrations (MIC) for all isolates was determined with ertapenem, imipenem and meropenem Epsilon tests (E-tests) (BioMèrieux, France). Briefly, a 0.5 McFarland inoculum of each isolate was lawned onto a Mueller-Hinton agar plate (Diagnostic Media Products, South Africa) and the E-test was placed at the center of the plate and incubated in ambient air at 35-37^0^C for 18-24 hours. The MIC was read at the point where the inhibition ellipse intersects with the test strip using the 2023 CLSI M-100 breakpoints [imipenem, meropenem, and doripenem resistance: ≥ 4 µg/mL; ertapenem resistance: ≥ 2 µg/mL]^12^.

The colistin MIC for the isolates were determined with broth microdilution (BMD). Colistin BMD was performed according to the Clinical and Laboratory Standards Institute (CLSI) document M07-A10 in an untreated 96-well microtiter polystyrene plate ^13^. Following subculture of the isolates on BAP’s, a 0.5 McFarland was inoculated into microtiter wells with colistin concentration ranging from 0.125 µg/mL to 64 µg/mL. The microtiter plates were incubated in ambient air at 35-37°C for 18-24 hours. The MIC was read at the first well with no macroscopically visible bacterial growth and interpreted using the 2023 CLSI M-100 breakpoints [Colistin resistance: ≥ 4 µg/mL] ^12^.

The modified carbapenem inactivation method (mCIM) was used to phenotypically screen the isolates for carbapenemase production ^14^.

### DNA extraction and PCR

A second subculture on BAP was done to obtain fresh cultures for DNA extraction and PCR to determine the presence of carbapenemase and colistin mobile resistance (*mcr*) genes. The isolates were further screened molecularly using multiplex PCR for *bla*_IMP_, *bla*_KPC_, *bla*_VIM,_ *bla*_OXA_, and *bla*_NDM_ carbapenemases as well as for *Mcr -1, Mcr -2, Mcr -3, Mcr-4,* and *Mcr-5* colistin resistance genes. The PCR screening included one multiplex PCR for *bla_VIM_*, *bla_OXA_* and *bla_NDM_* primers and two singleplex PCRs for both *bla*_KPC_ and *bla*_IMP_.The primers and the conditions used are shown in Table S1. In-house positive controls for carbapenemases were used while *mcr*-positive controls were provided by the Technical University of Denmark.

### Genomic and bioinformatic analyses

Owing to funding restrictions, we could only sequence 34 isolates. Therefore, we used the phenotypic data of the isolates to select the subset of 34 out of 40 isolates for genomic DNA extraction and sequencing; isolates with resistance to more than three antibiotics were prioritized for the whole-genome sequencing. gDNA were extracted from the isolates using the MagnaPure 96 instrument (Roche, South Africa) from 24-hour BAP cultures. The gDNA were sequenced on an Illumina Miseq at the genomic core sequencing facility of the National Institute of Communicable Diseases (NICD) (Johannesburg, South Africa). The generated fastQ and fastA files were submitted to GenBank under bioproject number PRJNA850834.

The genomes were annotated with NCBIs PGAP. The generated .gff files were used to delineate the genetic environment and associated mobile genetic environment of the resistance genes. The species of each of the isolates were confirmed using NCBIs ANI. ResFinder (https://cge.food.dtu.dk/services/ResFinder/) was used to determine the resistance determinants in the genomes. The clonality of the isolates and their multi-locus sequence typing (MLST) numbers were identified using MLST 2.0 (https://cge.food.dtu.dk/services/MLST/)^15^. The plasmid replicon genes or incompatibility groups and associated contigs were determined using PlasmidFinder (https://cge.food.dtu.dk/services/PlasmidFinder/). Mobile Element Finder was used to identify the mobile genetic elements (https://cge.food.dtu.dk/services/MobileElementFinder/) in the genomes, while the virulence genes were identified using VirulenceFinder (https://cge.food.dtu.dk/services/VirulenceFinder/).

#### Phylogenomics

The genomes of *K. pneumoniae, Enterobacter cloacae, Citrobacter portucalensis,* and *E. coli* isolates that are carbapenem- and/or colistin-resistant were curated from NCBI and PATRIC for purposes of determining their evolutionary relationships with this study’s isolates. These curated genomes were categorized geographically into South Africa, Africa, and global for the phylogenetic analyses. This study’s genomes and those from each of the three categories described above were aligned using Clustalw, which selected and aligned genes and sequences that were common to all aligned isolates. A minimum of 1000 genes from each isolates’ genome were thus aligned for the core genome phylogenetic tree construction. The phylogenies were subsequently inferred using RAxML (using a bootstrap reassessment of 1,000 x) and annotated with Figtree.

The ARGs from this study’s genomes and those of the genomes curated from NCBI and PATRIC were obtained from NCBI or ResFinder (https://cge.food.dtu.dk/services/ResFinder/). These ARGs were tabulated and arranged according to the host isolates’ location on the phylogenetic tree to provide a comparative phylogenomic view of the ARGs per clone or evolutionary distance.

### Ethical approval

This research was approved by the Ethical Review Board of the School of Medicine, University of Pretoria, South Africa.

## Results

### Outbreak settings and demographics

An increase in carbapenem-resistant infections were observed between September 2019 and March 2020 within the neonatal unit 4 at Tembisa hospital, South Africa. This outbreak involved 45 neonates who were born between August 2019 and March 2020 (Table S2). The affected infants were aged between 1 and 70 days (Table S2). Blood cultures (n = 26 specimen), and rectal swabs (n = 23 specimen) formed the most common clinical specimens collected from the infants. Single specimens were taken from tracheal aspirate (n = 1), urine (n = 1) and pus swab (n =1). Some of the specimens were taken from the same patient/infant (duplicates), hence the higher number of specimens than that of infants.

### Initial identification and antibiotic sensitivity testing

The isolates from the specimen were initially identified by Vitek 2 as *K. pneumoniae,* with seven specimens having a mixture of both *K. pneumoniae* and *Citrobacter* spp. (n = 2), *Enterobacter* spp. (n = 2), and/or *E. coli* (n = 3). Vitek II susceptibility testing found two isolates to be resistant to colistin with MICs ≥ 64 mg/uL and 19 isolates to be resistant to at least one carbapenem with an MIC of ≥ 2 mg/uL (Table S2).

*Carbapenemase production, MICs, and PCRs.* The results of mCIM, carbapenems MICs (E-test except for ertapenem for which Vitek 2 was used), colistin BMD, and PCRs are summarized in Table 1. All except four of the isolates were resistant to at least one carbapenem (imipenem and/or ertapenem) from the E-test; yet the four non-resistant isolates harboured a *bla*_OXA-48_-like carbapenemase. There were 31 imipenem- or ertapenem-resistant isolates, and 27 meropenem-resistant isolates while 27 isolates were resistant to all three carbapenems. All except seven isolates were colistin-resistant; these seven resistant isolates had no *mcr-1* to *-5* genes (Table 1).

**Table 1.**
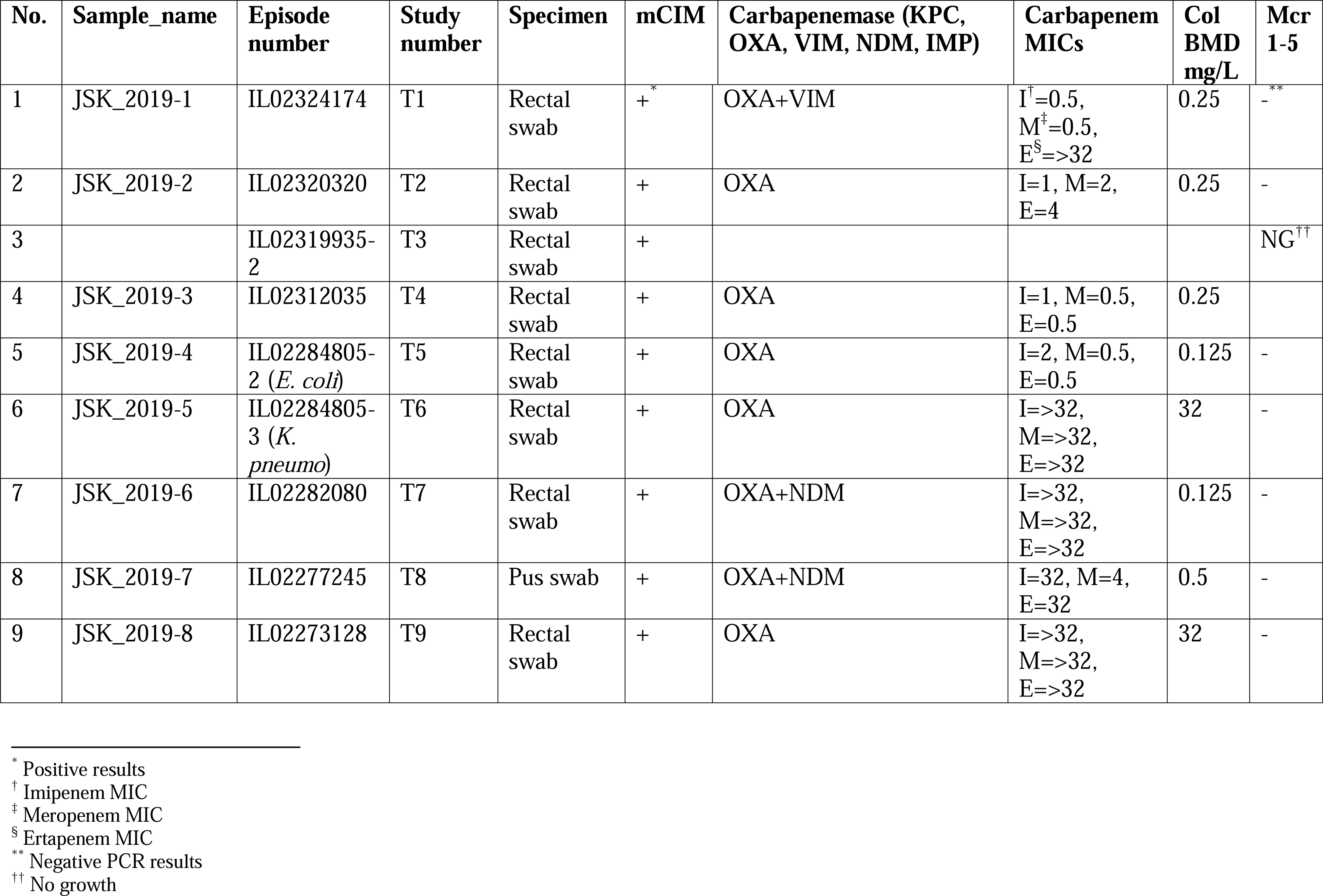
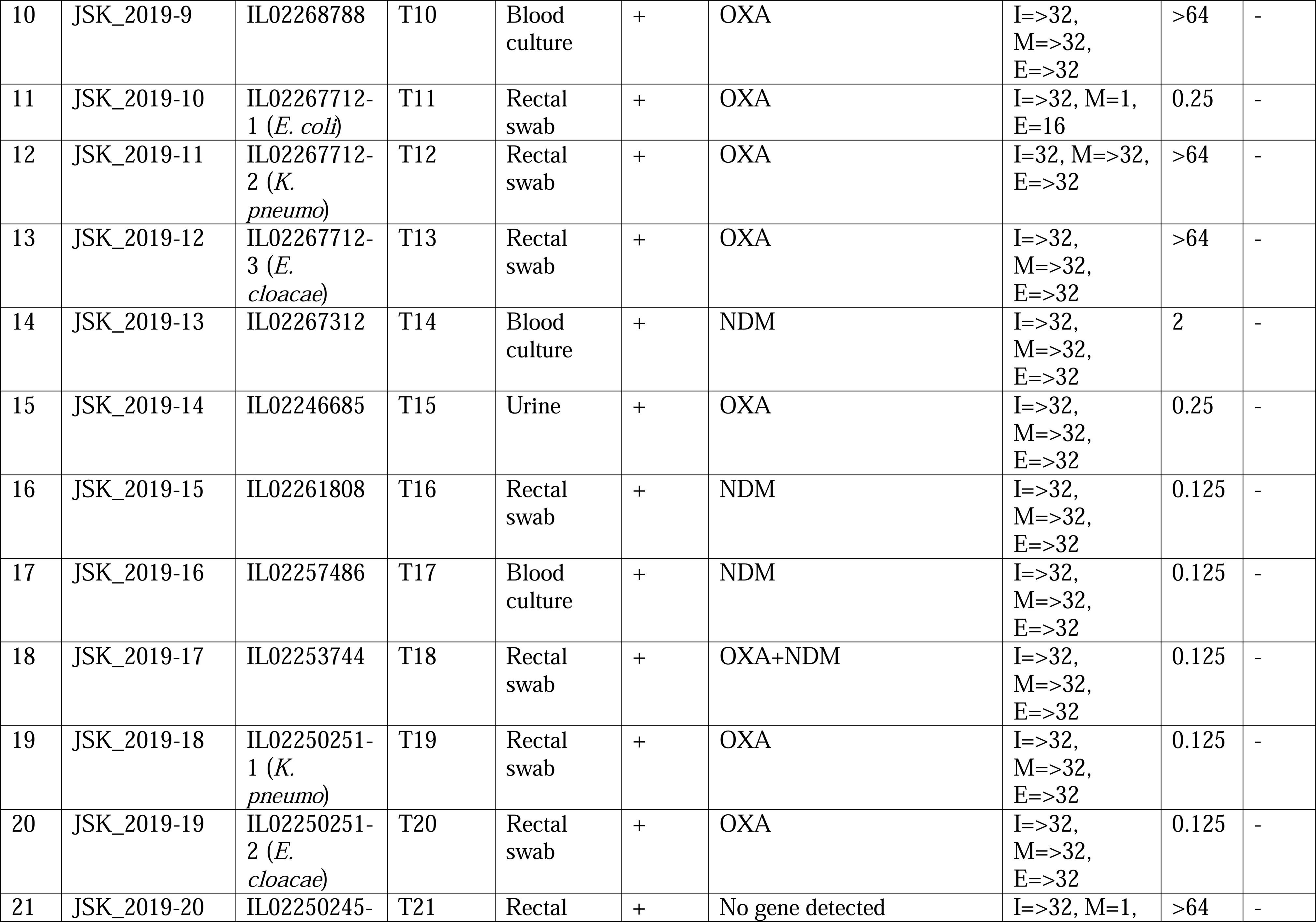
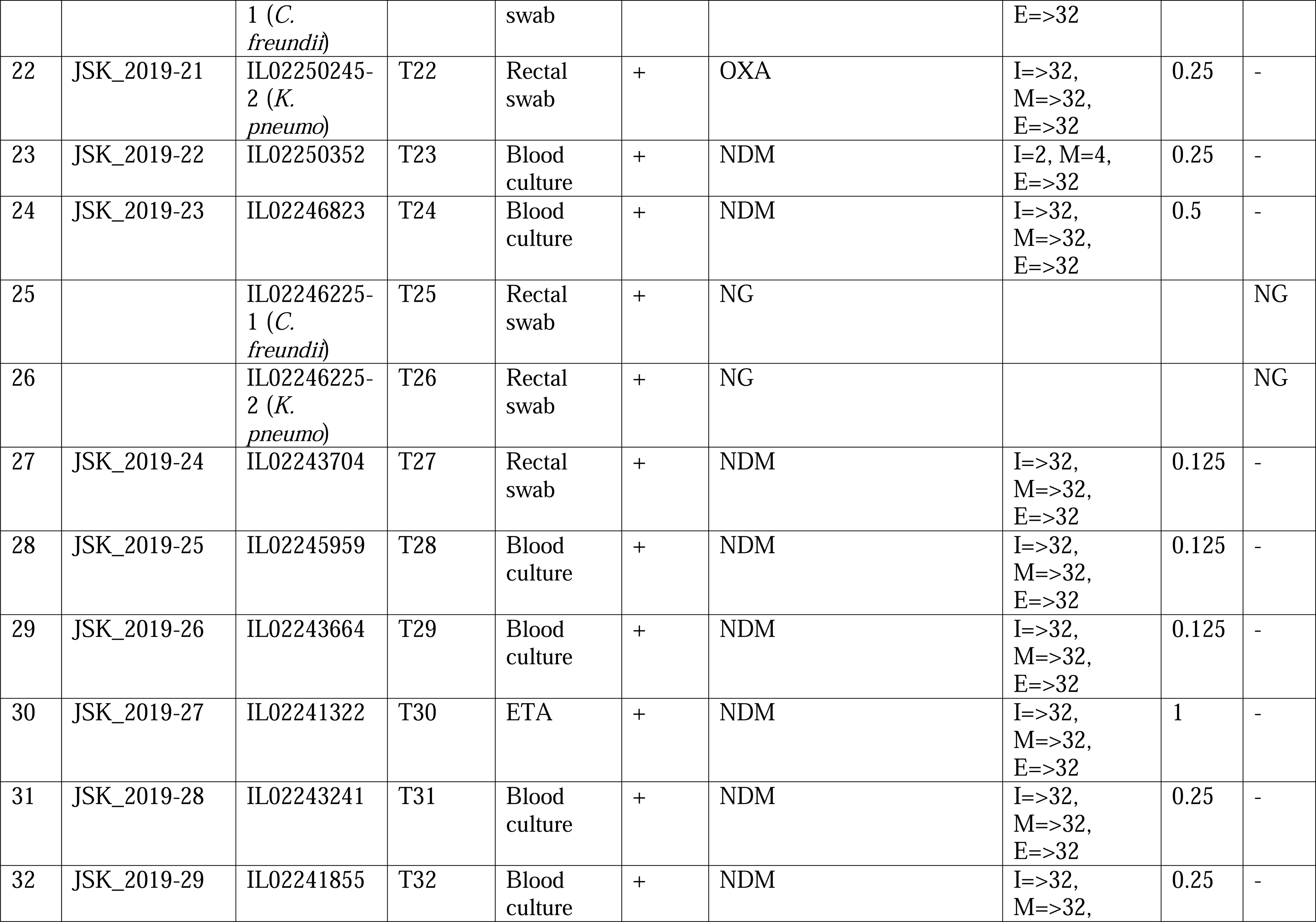
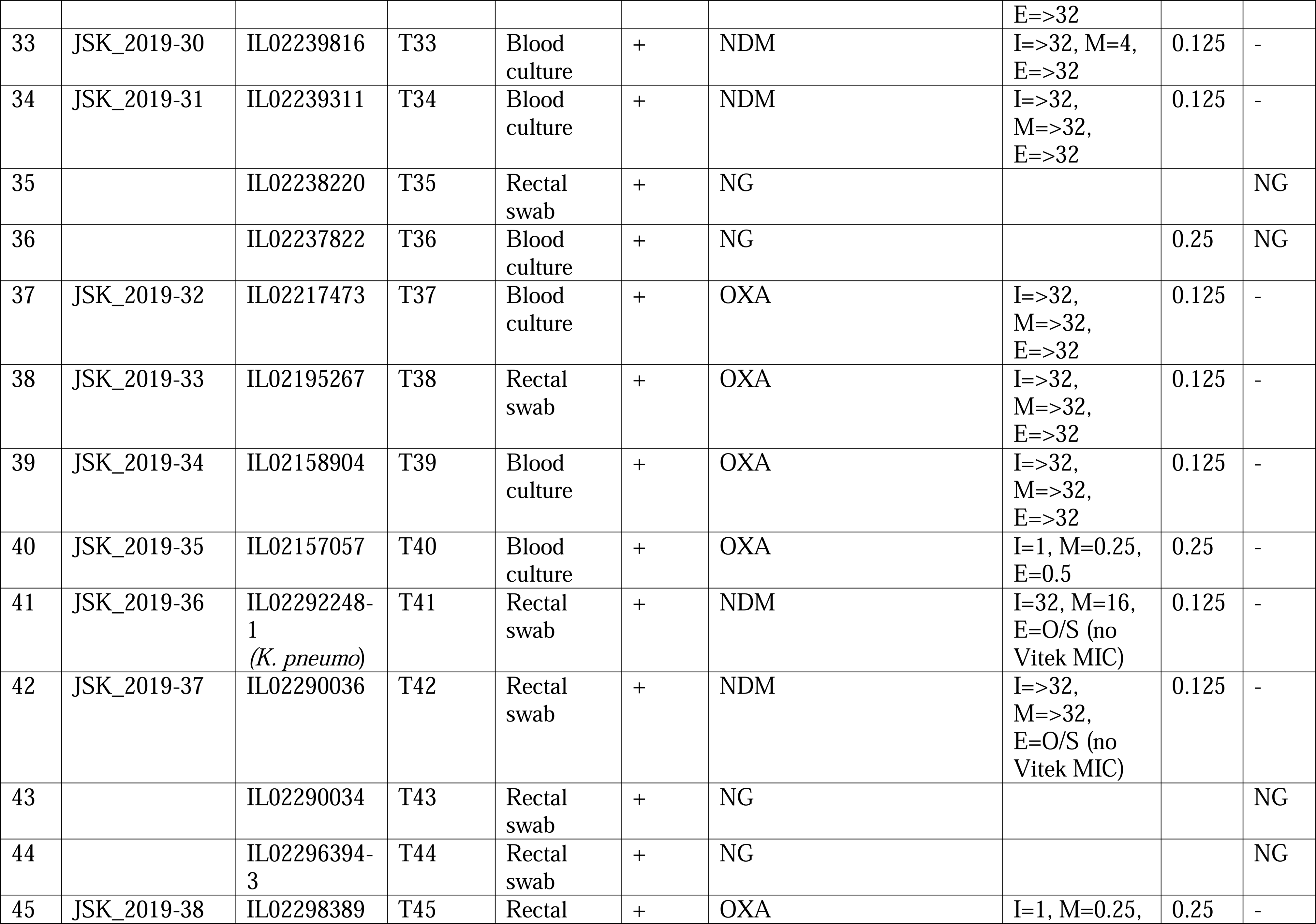
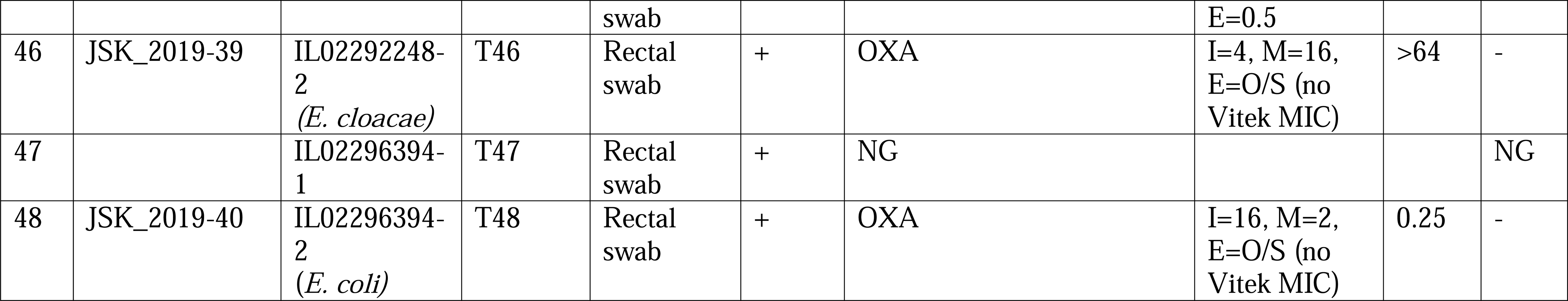
Summary of phenotypic, biochemical, and molecular (PCR) results for the carbapenem-resistant strains.

All the isolates were positive for the mCIM and PCR carbapenemase tests, except T21 (JSK_2019-20), which was mCIM-positive, but PCR-negative. There were 15 NDM-positive isolates, 18 OXA-48–like–positive isolates, three NDM + OXA-48–like–positive isolates, and one VIM + OXA-48–like–positive isolates. None of the isolates was *mcr-1* to -5–positive. Compared with OXA-48–like–positive isolates, all NDM-positive isolates were resistant to all three carbapenems except T33 (meropenem MIC = 4 µg/mL), T23 (imipenem MIC = 24 µg/mL; meropenem MIC = 4 µg/mL), and T8 (OXA-48–like and NDM-positive; meropenem MIC = 4 µg/mL); see Table 1.

#### Genomic analyses

### Identification and resistance profiles

The species of the isolates were confirmed by their average nucleotide identity (ANI) on NCBI. Of the 34 sequenced isolates, 27 were *Klebsiella pneumoniae,* three were *E. coli* (T-5, T-11, and T-48), three were *E. cloacae* (T-13, T-20, and T-46), and one was *Citrobacter portucalensis* (T-22). The inferred resistance profiles of the isolates, from their resistance genes, showed that 33/34 isolates were multidrug resistant. Except *E. cloacae* (T13, T20, and T46) and T48 (*E. coli*), all the other isolates were resistant to at least four antibiotic classes: aminocyclitol, aminoglycosides, amphenicol, βeta-lactam, folate pathway antagonist, fosfomycin, macrolide, peroxide, quaternary ammonium compound, quinolone, rifamycin, and tetracycline (Table S3). Only one isolate had a tetracycline resistance gene.

### Resistance determinants

The distribution of the antibiotic resistance genes per isolate is shown in Table S3, with the most frequently identified being *oqxAB* (n = 28 isolates), *fosA* (n = 27 isolates), *sul-1* & *sul-2* (n = 23 isolates), *qacE* (n = 22 isolates), *mphA* (n = 20 isolates), *aac(6’)-Ib-cr* (n = 20 isolates), *aph(3”)-Ib* (n = 17 isolates), *aph(6)-Id* (n = 17 isolates), *aadA16* (n = 16 isolates)*, dfrA27* (n = 16 isolates), *arr-3* (n = 16 isolates), *bla*_CTX-M-15_ (n = 16 isolates), *bla*_TEM-1B_ (n = 16 isolates), *bla*_OXA-48_ (n = 15 isolates), *bla*_SHV-187_ (n = 14 isolates), and *rmtC* (n = 12 isolates). *bla*_NDM-1_ was present in 13 isolates (Table S3).

Common mutation-based resistance mechanisms were found in *gyrA* (n = 25 isolates), *ompK36* (n = 26 isolates), and *ompK37* (n = 26 isolates). Nineteen *K. pneumoniae* isolates viz., T8, T9, T10, T12, T14, T16, T17, T18, T19, T23, T27, T28, T29, T30, T31, T32, T33, T37, and T39, had between 15 and 22 resistance genes (Table S3).

Discrepancies between the PCR data and the whole-genome sequencing (WGS) data were observed. PCR did not identify *bla*_OXA-181_ in strain T21 (JSK-2019-20), albeit WGS identified this gene and mCIM showed the presence of a carbapenemase. Moreover, the WGS data did not confirm the presence of VIM + OXA-48, NDM + OXA-48, and the presence of OXA and NDM in some of the isolates. Indeed, some of the isolates could not be revived for WGS after the PCR screening step. Evidently, some might have lost their plasmids during the subsequent culturing steps to obtain 24-hour genomic DNA for the WGS. However, the presence of both NDM and OXA-48/181 in the isolates were mostly confirmed by both PCR and WGS (Tables 1 and S3).

### Virulence factors

Among the 34 sequenced isolates, 24 virulence genes were identified: *air* (n = 1 isolate), *ccl* (n = 12 isolates), *cia* (n = 2 isolates), *chuA* (n = 2 isolates), *cvaC* (n = 1 isolate), *eilA* (n = 2 isolates), *etsC* (n = 1 isolate), *fyuA* (n = 18 isolates), *gad* (n = 2 isolates), *hylF* (n = 1 isolate), *KpsMII_K5* (n = 1 isolate), *KpsE* (n = 1 isolate), *ipfA* (n = 2 isolates), *iroN* (n = 1 isolates), *irp2* (n = 18 isolates), *iss* (n = 2 isolates), *iucC* (n = 1 isolate), *iutA* (n = 28 isolates), *mchF* (n = 1 isolate), *ompT* (n = 2 isolates), *sitA* (n = 1 isolate), *terC* (n = 12 isolates), *traT* (n = 5 isolates), and *yfcV* (n = 1 isolate). *E. coli* isolates T11 and T48 had the highest number of virulence genes (n = 17 and 13, respectively), while the other isolates harboured between one and four genes.

### Plasmid replicon/incompatibility types

An average of four plasmid incompatibility groups were found in each isolate, with higher numbers being detected in isolates T1 (n = 7), T2 (n = 9), T11 (n = 5), T18 (n = 8), T20 (n = 5), T22 (n = 6), T39 (n = 6), and T40 (n = 6). The remaining isolates had between one and four incompatibility groups, with four plasmid types per isolate being very dominant. There were 25 plasmid types, of which IncFIA(HI1) (n = 5 plasmid types), Col(pHAD28) (n = 8 plasmid types), ColKP3 (n = 10 plasmid types), ColRNAI (n = 14 plasmid types), IncFIB(K) (n = 15 plasmid types), IncFIB(pNDM-Mar) (n = 7 plasmid types), IncFII(Yp) (n = 16 plasmid types), IncL (n = 16 plasmid types) and IncX3 (n = 10 plasmid types) were most common.

Of the 55 mobile genetic elements (MGEs) identified in the isolates, *IS*Vsa3 (n = 5 isolates), *IS*Ec15 (n = 7 isolates), *IS*Ec36 (n = 10 isolates), *IS*Kpn19 (n = 11 isolates), *IS*Kpn21 (n = 11 isolates), *IS*Kpn26 (n = 11 isolates), *IS*Kpn28 (n = 11 isolates), *IS*Kpn34 (n = 11 isolates), *IS*Kpn43 (n = 11 isolates), *IS*Sen4 (n = 11 isolates), *IS*26 (n = 12 isolates), *IS*Ec33 (n = 13 isolates), *IS*Kox1 (n = 13 isolates), *IS*5075 (n = 19 isolates), *IS*6100 (n = 22 isolates), *IS*Kox3 (n = 24 isolates), and *IS*Kpn1 (n = 20 isolates) were prominent. The number of MGEs per isolate ranged from three to 16, with an average of nine MGEs per isolate.

The MGEs associated with *bla*_NDM-1_ and *bla*_OXA-48/-181_ are shown in Figure 1. Irrespective of the strain (MLST) or species, the same genetic environment in the same or reversed orientation, or a truncated version of it (in the case of OXA-181), was found within the immediate flanks of *bla*_NDM-1_, *bla*_OXA-48_, and *bla*_OXA-181_ (Fig. 1).

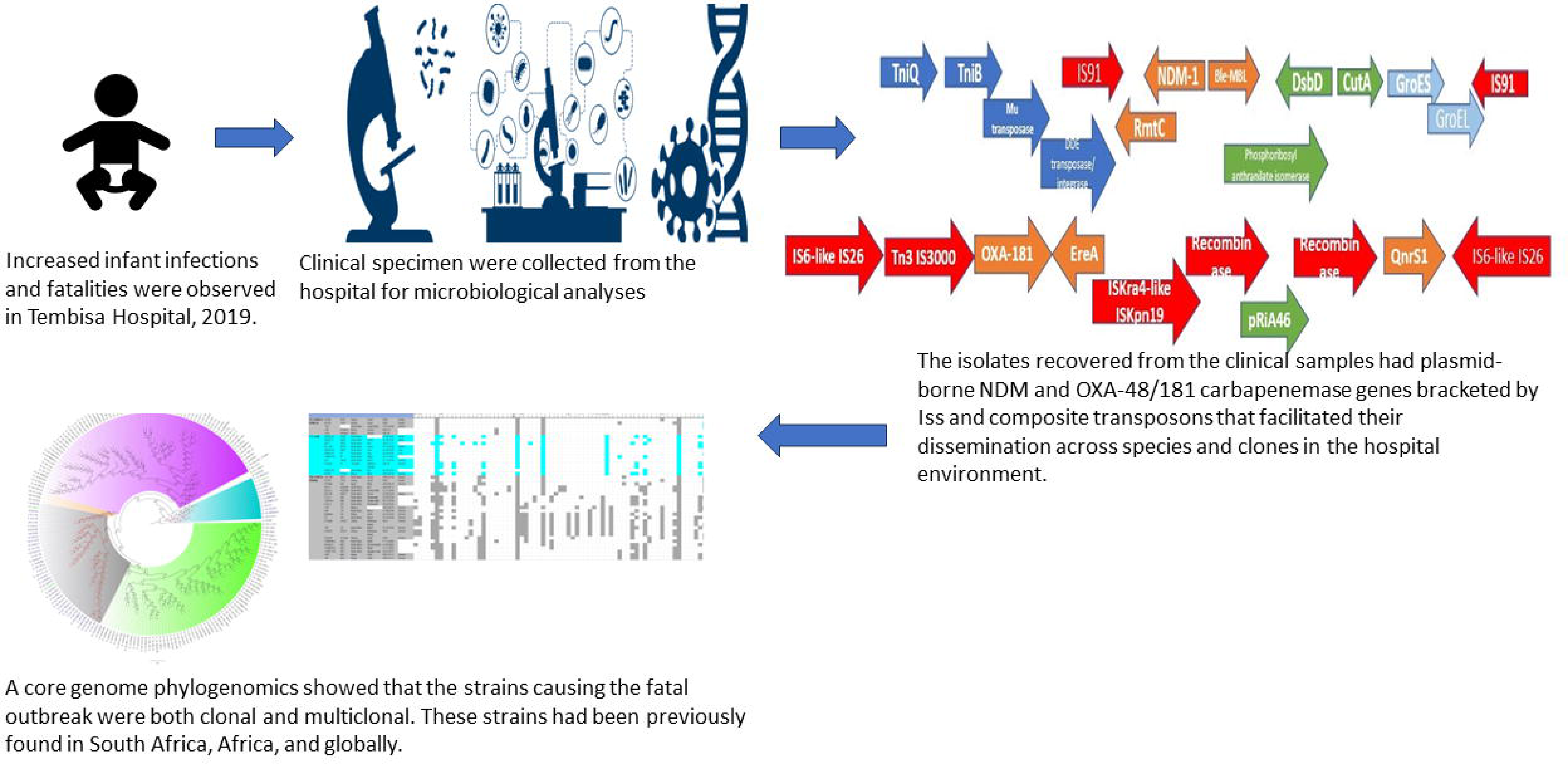

The clonality of the isolates was determined using their MLST (see Tables 1 & S3). All the three *E. coli* strains were not related and three of the *K. pneumoniae* strains (i.e., ST5785, ST25, and ST297) were also singletons with no relation with the other *K. pneumoniae* strains that occurred in multiples: ST17 (n = 2); ST1266 (n = 3); ST307 (n = 9); ST152 (n = 13). Whereas the single *K. pneumoniae, E. coli,* and *C. portucalensis* clones as well as the three *E. cloacae* clones were isolated in 2020 from rectal swabs, *K. pneumoniae* ST307 and ST152 were identified between 2019 and 2020 from different specimen types: pus, ETA, rectal swab, blood culture, and urine.

*K. pneumoniae* ST17 was only isolated in 2019 from blood cultures only.

A uniformity in resistance profiles, ARGs, virulence genes, plasmid types, and MGEs was not observed among isolates of the same clone although some genes and genetic elements were uniformly present in members of the same clone (Table S3).

#### Phylogenomics

The evolutionary relationships and antibiotic resistance patterns between this study’s isolates and other closely related isolates are depicted in Figures 2-8, Figures S1-S9, and Table S4. With some exceptions, isolates belonging to the same MLST (clones) were found to be closely related (found on the same branch with bootstrap values of more than 50) to each other than those from different MLST clones on the trees. Moreover, the resistance profiles of the clones/isolates found on the same branch (with very close evolutionary distance) were very similar, albeit some differences were also observed. For instance, irrespective of country, source, and date of isolation, *E. coli* ST131 clones had very similar ARG profiles (Fig. 2), suggesting a possible international transmission of that clone with their associated ARGs. Similar trends were also observed in closely related *K. pneumoniae* clones and isolates on the same cluster or branch. Notably, *E. cloacae* isolates that were of very close evolutionary distance in Fig. 7 had very similar resistance profiles, which differed from those that were distant and found on different branches and clusters. These uniform patterns were observed in strains that were isolated from different parts of the world and from different clinical and environmental specimen types.

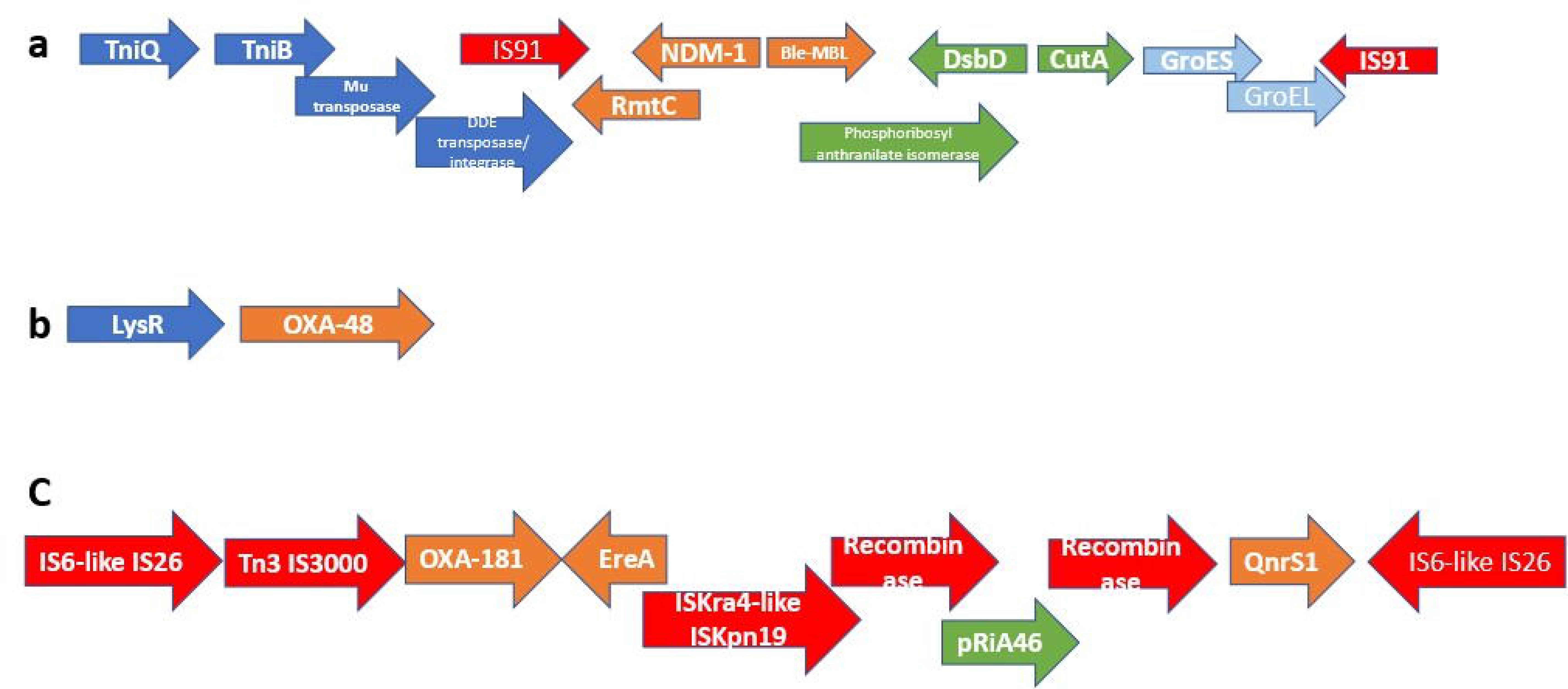

## Discussion

In this study, we describe an outbreak of fatal carbapenem-resistant *K. pneumoniae, E. coli,* and *E. cloacae* infections in a tertiary hospital in Gauteng, South Africa, which contributed to the mortality of at least 10 neonates. It is worrying to note that the strains were mostly multidrug-resistant, with all except four of the strains being resistant to at least four antibiotic classes (Figures 2-8 and S1-S9; Tables 1 and S1-S4). More concerning is the presence of carbapenemases in all the strains, which confer resistance to almost all β-lactams. Although no *mcr* genes were identified, seven strains were resistant to colistin, a last-resort antibiotic for carbapenem-resistant Gram-negative bacterial infections ^7^.

Outbreaks involving *K. pneumoniae* and *E. coli* in hospital settings are common; including fatal outbreaks in neonatal units as reported in this study ^3^. Indeed, this is not the first outbreak to occur in South Africa as well. However, this study presents one of the most comprehensive molecular and genomic analyses of infection outbreaks involving these species and *E. cloacae* ^5^.

Although all the isolates were mCIM-positive and all-but-one was PCR-positive for carbapenemases, they were not all phenotypically resistant to carbapenems; at least four isolates were carbapenem-susceptible (Table 1). This observation is not singular as some isolates with carbapenemases have been reported to be phenotypically susceptible to carbapenems. For such observations, it was concluded that the carbapenemases were not expressed, although they were present ^16,17^. For the PCR-negative but mCIM-positive isolate, there could be the presence of either a novel carbapenemase or a known carbapenemase that was not targeted by the PCR. The latter possibility was invalidated by the presence of *bla*_OXA-181_, which was identified by the whole-genome data but was not identified by the PCR. Thus, the mCIM agreed with the whole-genome sequencing data.

However, it is worth noting that carbapenem resistance is not only mediated by carbapenemases as other determinants such as hyperexpression of AmpCs, ESBLS, and efflux pumps and hypo-expression of porins also determine carbapenem resistance ^18–20^. The presence of a resistance gene with a non-commensurate phenotypic expression of the resistance gene is not limited to carbapenems alone. This is one of the reasons why molecular diagnostic tests for antibiotic resistance genes cannot be used solely as representative or confirmatory of phenotypically resistant pathogens. In this study, we also observed that the colistin-resistant isolates did not have *mcr* genes ^7,18^.

As recently observed in a molecular study to screen for carbapenemases and *mcr* genes, *mcr* genes are relatively less prevalent than carbapenemases ^21^. This phenomenon was also observed in this study as no *mcr* gene was identified, albeit seven isolates were colistin-resistant. This is a positive finding as colistin is currently the last-resort antibiotic for carbapenem-resistant infections ^7,22^. Moreover, the rarity of *mcr* genes will help reduce the spread of colistin resistance as *mcr* genes are mainly plasmid-borne and very mobile.

Similar findings with higher prevalence of *bla*_OXA-181/48_ and *bla*_NDM_ carbapenemase genes in Gram-negative bacteria has been reported in South Africa, confirming this observation, and showing that these two carbapenemases are common in South African hospitals ^16,23–27^. The phylogenomic analyses of the South African isolates confirm these observations and similar patterns (Table S4). Furthermore, the presence of other clinically important resistance genes such as *bla*_CTX-M-15_, *bla*_SHV_, *bla*_TEM_, *qnrB/S, oqxA/B, dfrA, Sul1/2/3, aac(6’)-lb-cr, aadA,* and *mph(A)* explains the multidrug and pan-drug resistance nature of the isolates. This might explain the failure of the clinicians to treat these infections, leading to the mortality of 10 infants.

As shown in Fig. 1, the carbapenemases in the isolates were flanked by other resistance genes and mobile genetic elements such as composite transposons: *IS*91 (*bla*_NDM_) and *IS*6-like *IS*26 (*bla*_OXA-181_). Instructively, irrespective of the strain in which any of the three carbapenemase i.e., *bla*_OXA-48_, *bla*_OXA-181_, and *bla*_NDM-1_, was found, their genetic support/environment were all the same. This similarity in synteny and flanks around these resistance genes is further corroborated by the fact that most of these isolates, irrespective of the species or the clone, had the same insertion sequences: *IS*26 (n = 12); *IS*Ec33 (n = 13); *IS*Kpn26 (n = 16); *IS*Sen4 (n = 16); *IS*5075 (n = 19); *IS*Kpn1 (n = 20); *IS*6100 (n = 22); *IS*Kox3 (n = 24) (Table S3: MGEs). Furthermore, each of the isolate had an average of three plasmid replicons/incompatibilities, with the least being one and the highest being nine. The commonest of these plasmid types among all the isolates were IncL (n = 16), IncFII(Yp) (n = 16), IncFIB(K) (n = 15), ColRNAI (n = 14), IncX3 (n = 10), and ColKP3 (n = 10). Owing to the break-up of the genomes into several contigs, it is difficult to associate these plasmid types with each ARG. However, taken together, these show that the ARGs in these MDR isolates are being disseminated through these MGEs (Table S3: Plasmid incompatibility).

These genetic environments and ARG flanks are also commonly observed in other studies involving the same or different species within Enterobacterales from both South Africa and globally, corroborating the role of MGEs in the transmission of carbapenemases and other ARGs within and across clones and species ^2,5,7,16,28^.

Although there are differences in the ARGs present in the isolates (Table S3: ARGs), there are very close similarities and uniformity in the ARGs present in the isolates. This uniformity in ARGs, MGEs, and plasmid incompatibility/replicon types across the isolates suggest that irrespective of the clones and species, there are plasmid-borne MGEs that are shuttling the ARGs between and across the species and clones ^5,7,16,28,29^. Therefore, there is both clonal and plasmid-mediated transmission of the same ARGs in the hospital, particularly when the outbreak occurred in two major wards: ward 4 and 4a-NICU.

In fact, this observation is also magnified in Figures 2-8, where the same ARGs were seen among isolates of the same clone and isolates with very close evolutionary distance. In particular, the phylogenomic analyses of the isolates show that the same ARGs, possibly hosted on MGEs, and clones are being disseminated across South Africa, Africa, and globally. It is observed from Figures 2-8 that the same clones among the three species cluster together closely in the phylogenetic tree alongside other clones. The presence of other clones clustering alongside the same clones is not surprising given the higher resolution of whole-genome-based typing compared to multi-locus sequence typing (MLST) that only uses seven house-keeping genes.

The *E. coli* isolates from this outbreak were of the same clone as *E. coli* clones ST58, ST69, ST87, ST103, ST131, ST457, ST616, and ST648, which were common among other isolates from South Africa, Africa, and globally. The resistance profiles within each of these clones or phylogenetic cluster were mostly uniform with multidrug resistance (Fig. 2-3).

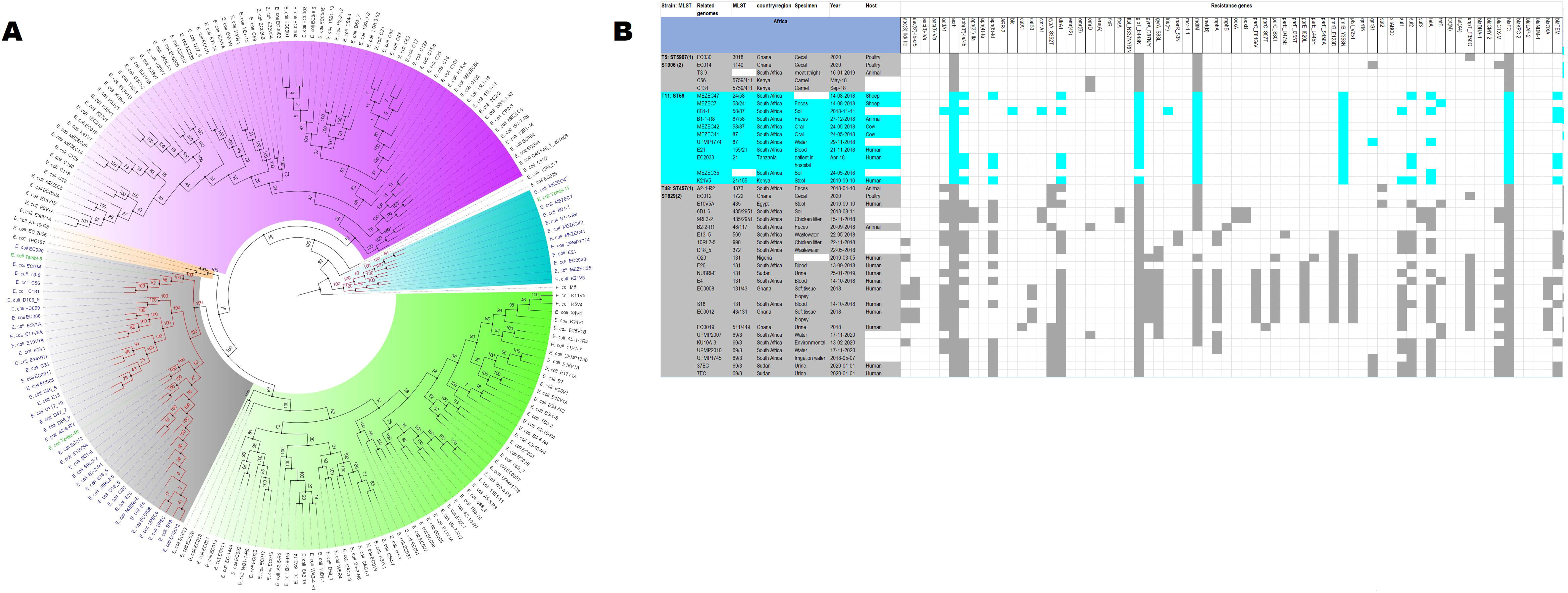

The *K. pneumoniae* clones in this study shared the same clone and phylogenetic cluster as *K. pneumoniae* ST14, ST15, ST17, ST25, ST101, ST152, ST231, and ST307, and the resistance profiles within each of these clonal clusters were very similar with multiple resistance genes (Fig. 4-6). Likewise, the *E. cloacae* isolates clustered with *E. cloacae* ST84 and ST456 with very similar multi-resistance profiles (Fig. 7-8). These observations support the fact that there is clonal and MGE-mediated transmission of these ARGs across the globe, which caused the outbreak in the hospital under study.

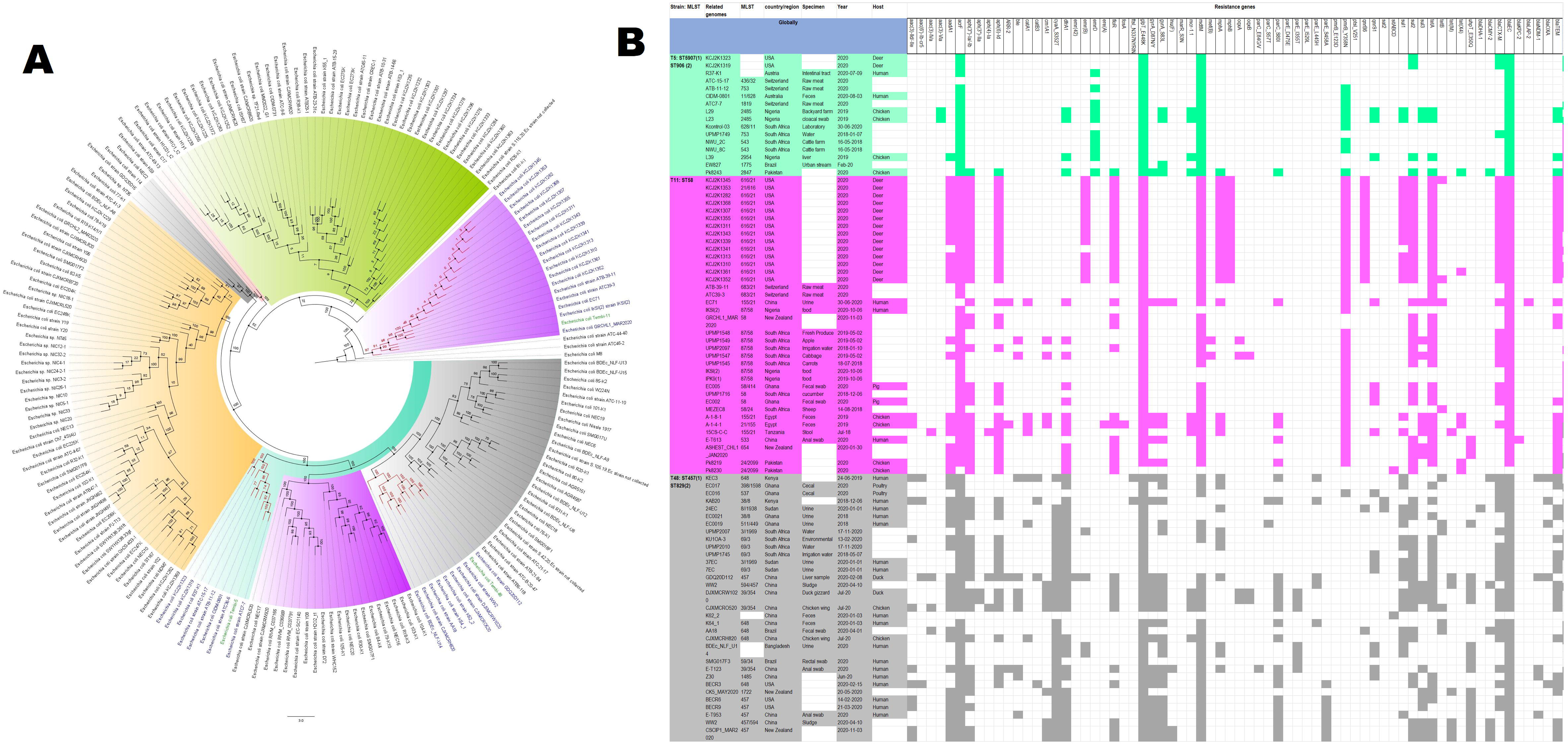

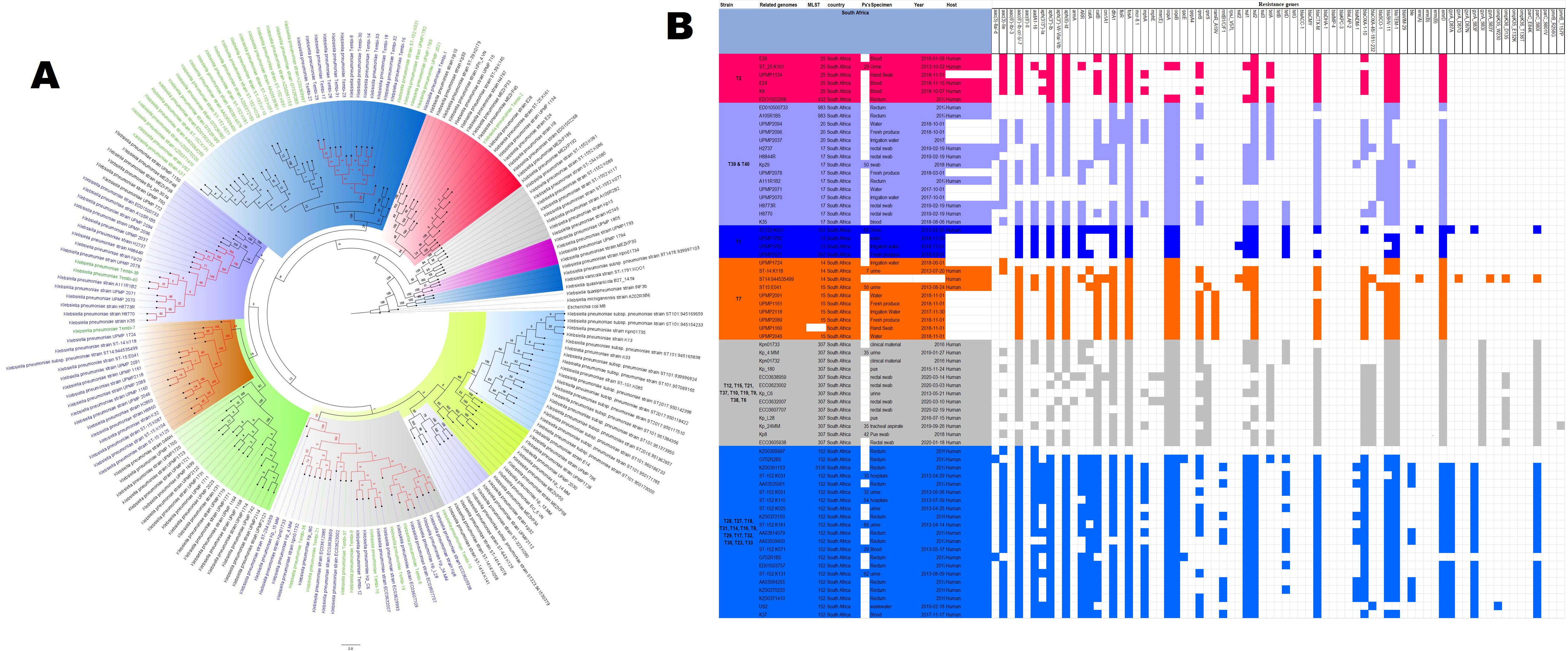

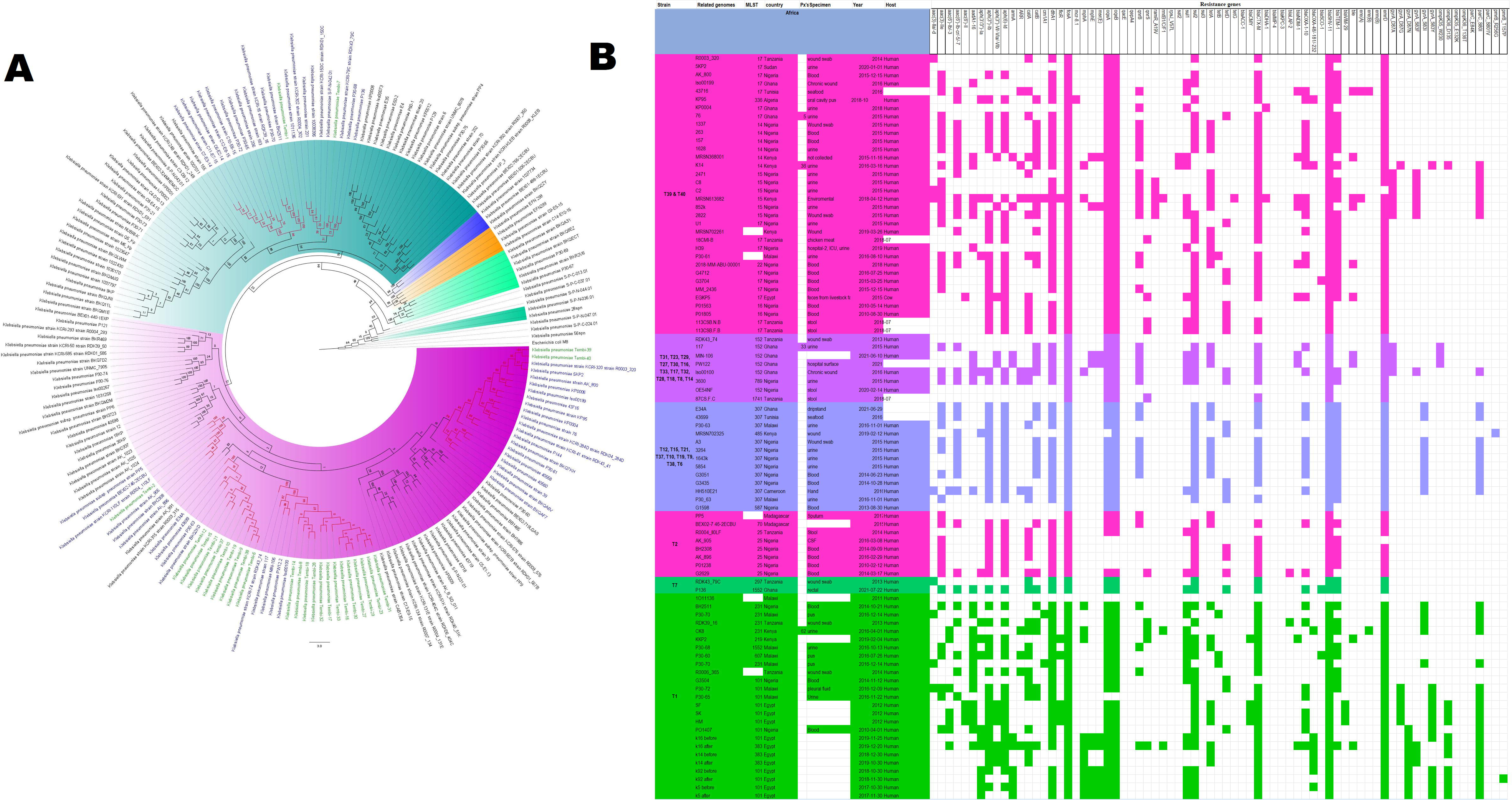

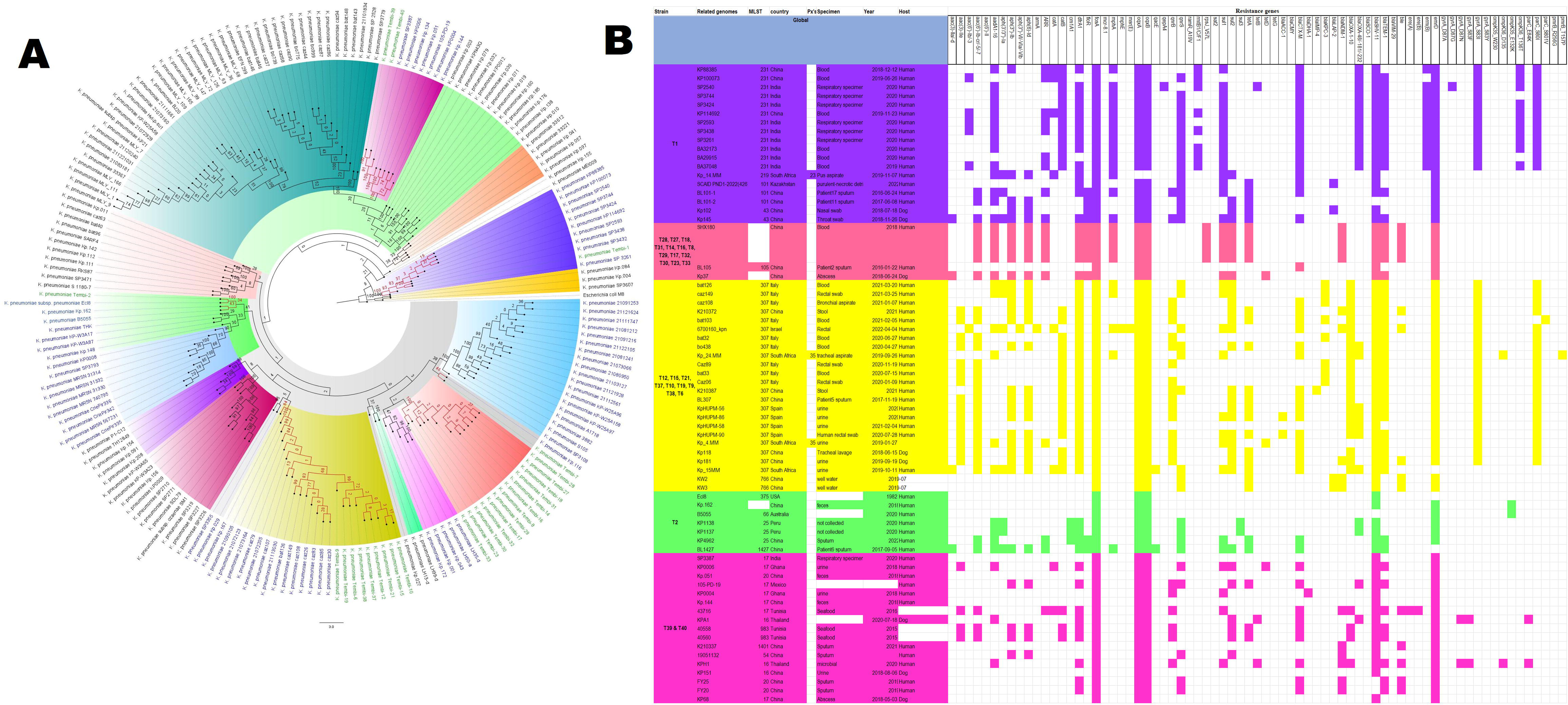
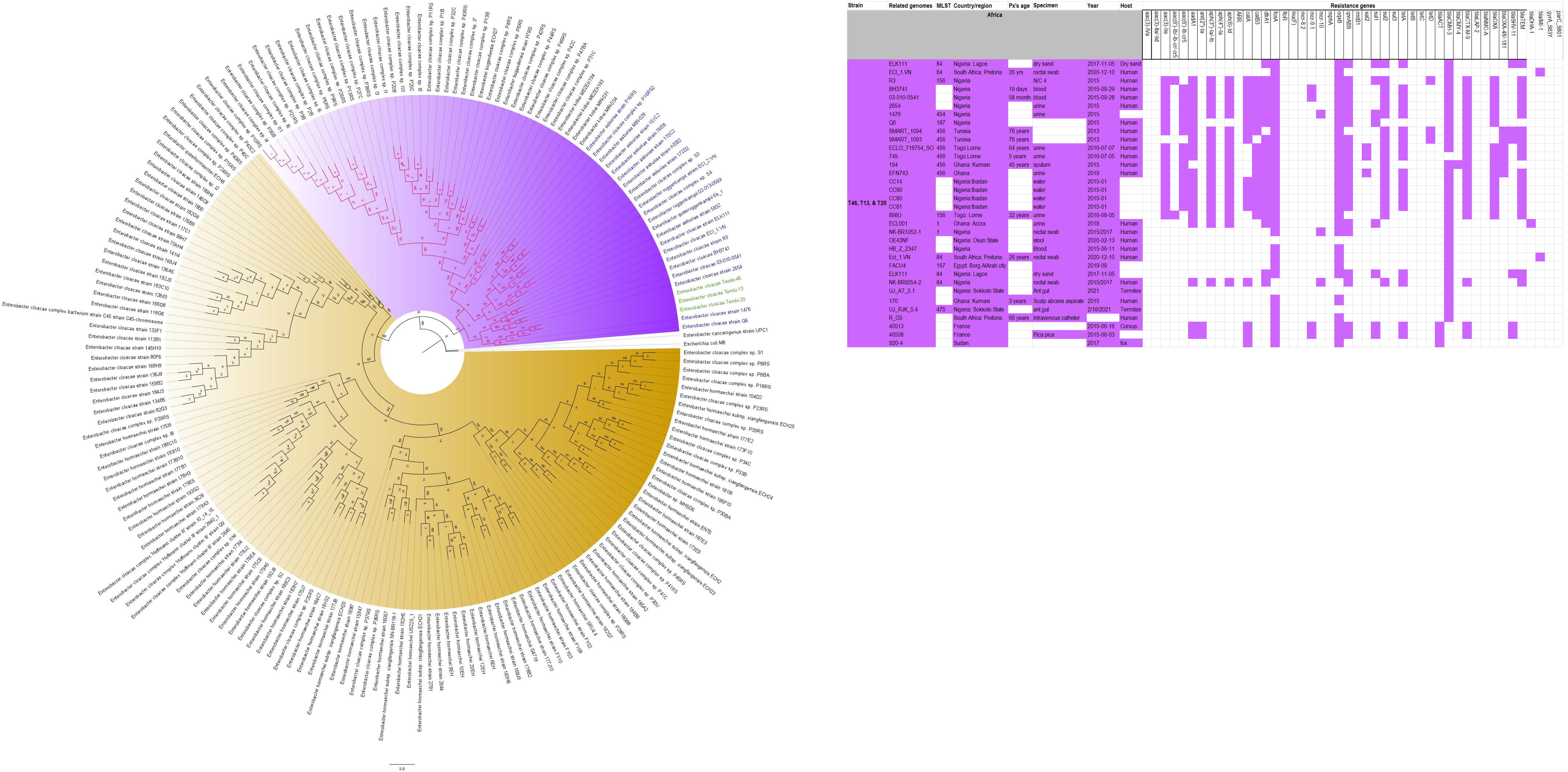

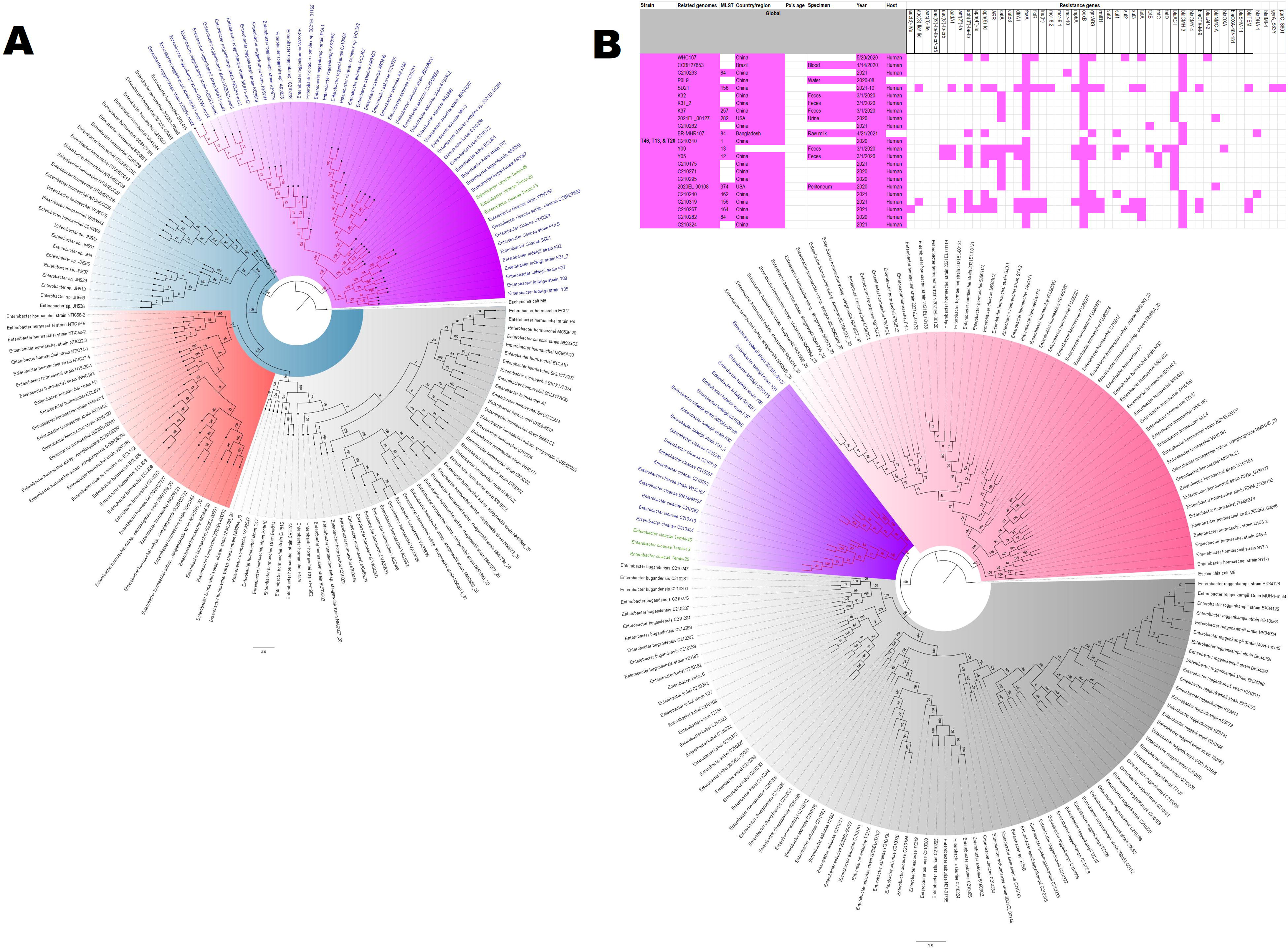

In summary, the outbreak that occurred at the Tembisa hospital was clonally and horizontally mediated, spreading multidrug-resistant infections and causing fatalities.

## Supporting information

Table S1. Primers, PCR conditions, & PCR Results.

Table S2. De-identified demographic data of the human hosts from which the strains were isolated.

Table S3. Raw sequence and genomic data of the strains

Table S4. Phylogenomic and comparative genomic data of the strains.

## Acknowledgement

We are grateful to the technical staff of the National Health Laboratory Service (NHLS), Tshwane Academic Division, University of Pretoria, for their assistance with the initial microbiological processing of the clinical samples.

## Author contributions

SD assisted with data capturing and data analysis; NEM assisted with data capturing and manuscript editing; AB and MS undertook phenotypic AST testing, data capturing, analysis of AST results, and reviewing of manuscript. RV assisted with initial data and isolates collation, and review of manuscript. MM performed laboratory work and reviewing of manuscript. NMM initiated the concept and design as well as assisted with the co-ordination of the project. JOS designed, supervised, and coordinated the execution of the project, undertook bioinformatics analyses and image generation, wrote the manuscript, and validated the final version of the manuscript for submission.

## Transparency declaration

The authors declare no conflict of interest and the funding agency (NHLS) had no role in the decision to write or submit this manuscript.

---

**Table S1.** Primers, PCR conditions, & PCR Results.

**Table S2.** De-identified demographic data of the human hosts from which the strains were isolated.

**Table S3.** Raw sequence and genomic data of the strains

**Table S4.** Phylogenomic and comparative genomic data of the strains.

**Figure S1. Comparative phylogenomics of *Escherichia coli* strains from both this study and other strains from South Africa**. The names of the strains from this study are coloured green. The names of closely related strains to this study’s strains are shown in blue. Branches holding this study’s strain with very high bootstrap values (>50%) are shown in red to depict isolates with very close evolutionary distance.

**Figure S2. Comparative phylogenomics of *Escherichia coli* strains from both this study and other strains from Africa**. The names of the strains from this study are coloured green. The names of closely related strains to this study’s strains are shown in blue. Branches holding this study’s strain with very high bootstrap values (>50%) are shown in red to depict isolates with very close evolutionary distance.

**Figure S3. Comparative phylogenomics of *Escherichia coli* strains from both this study and other strains from the world**. The names of the strains from this study are coloured green. The names of closely related strains to this study’s strains are shown in blue. Branches holding this study’s strain with very high bootstrap values (>50%) are shown in red to depict isolates with very close evolutionary distance.

**Figure S4. Comparative phylogenomics of *Klebsiella pneumoniae* strains from both this study and other strains from Africa**. The names of the strains from this study are coloured green. The names of closely related strains to this study’s strains are shown in blue. Branches holding this study’s strain with very high bootstrap values (>50%) are shown in red to depict isolates with very close evolutionary distance.

**Figure S5. Comparative phylogenomics of *Klebsiella pneumoniae* strains from both this study and other strains from the world**. The names of the strains from this study are coloured green. The names of closely related strains to this study’s strains are shown in blue. Branches holding this study’s strain with very high bootstrap values (>50%) are shown in red to depict isolates with very close evolutionary distance. Both S5A and S5B are phylogenetic analyses of global strains.

**Figure S6. Comparative phylogenomics of *Enterobacter cloacae* strains from both this study and other strains from Africa**. The names of the strains from this study are coloured green. The names of closely related strains to this study’s strains are shown in blue. Branches holding this study’s strain with very high bootstrap values (>50%) are shown in red to depict isolates with very close evolutionary distance.

**Figure S7. Comparative phylogenomics of *Citrobacter portucalensis* strains from both this study and other strains from the whole world**. The names of the strains from this study are coloured green. The names of closely related strains to this study’s strains are shown in blue. Branches holding this study’s strain with very high bootstrap values (>50%) are shown in red to depict isolates with very close evolutionary distance.

