## Supplementary figures and images for "Molecular Epidemiology of multidrug-resistant *Klebsiella pneumoniae, Enterobacter cloacae,* and *Escherichia coli* outbreak among neonates in Tembisa Hospital, South Africa"

### Fig. S1

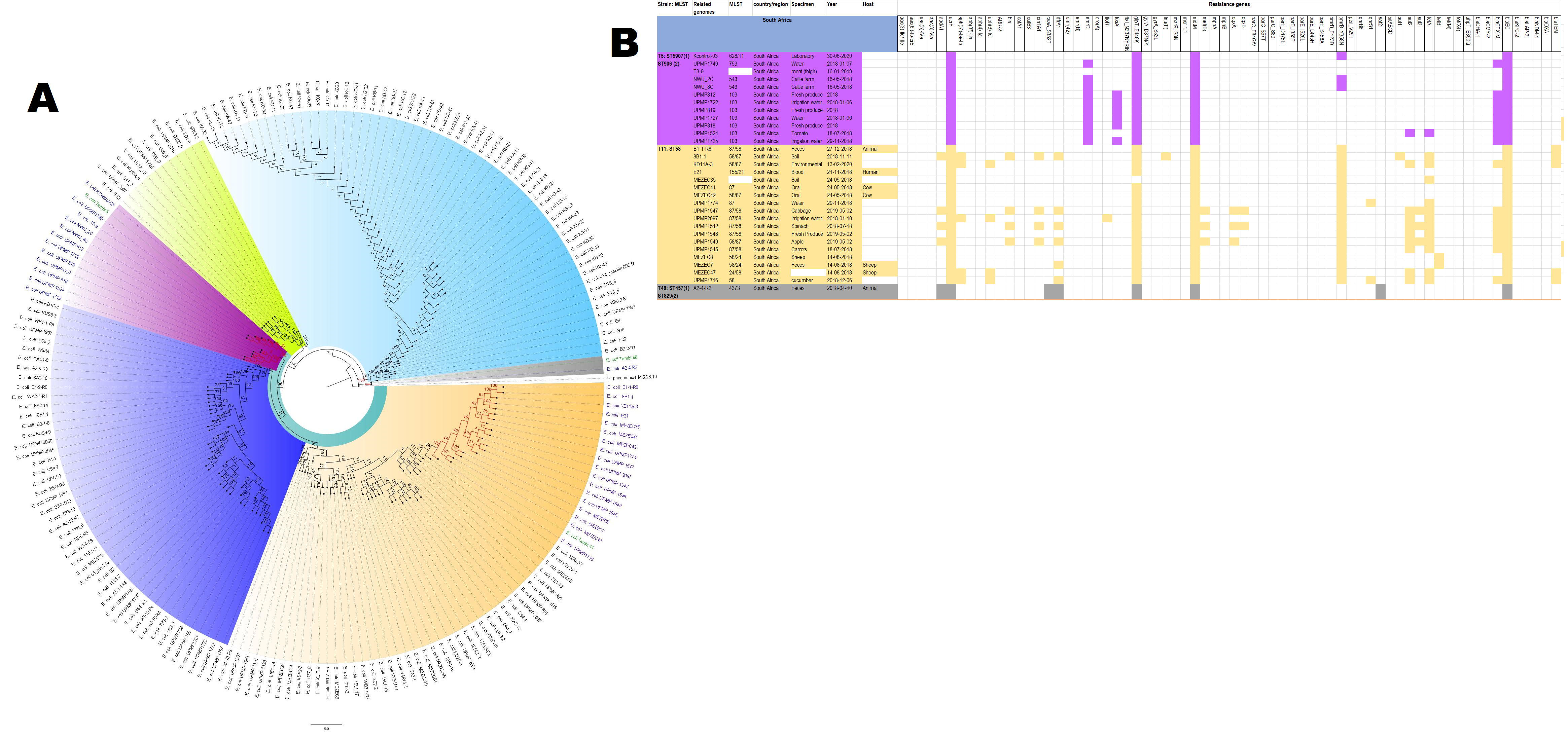

### Fig. S2

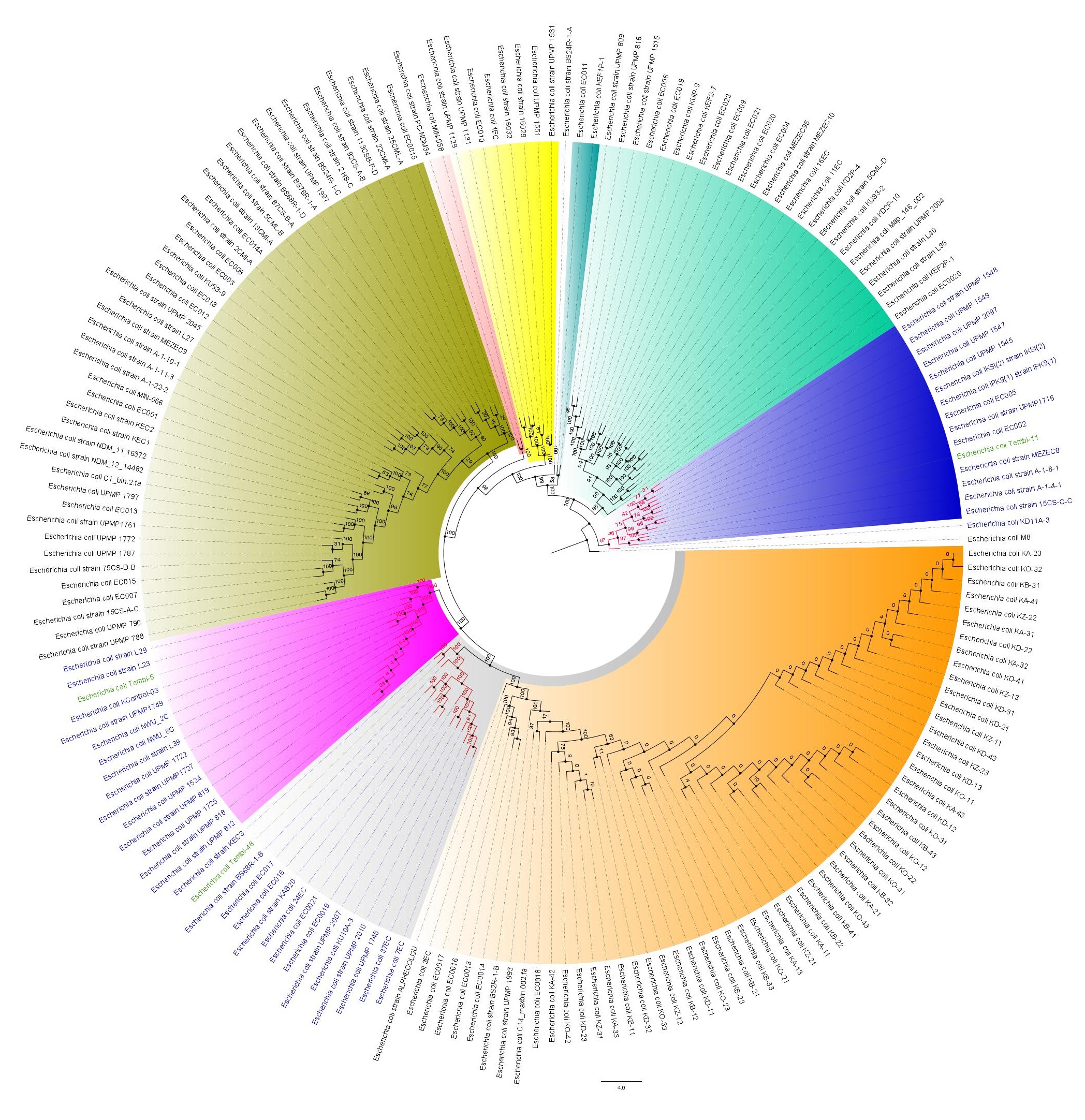

### Fig. S3

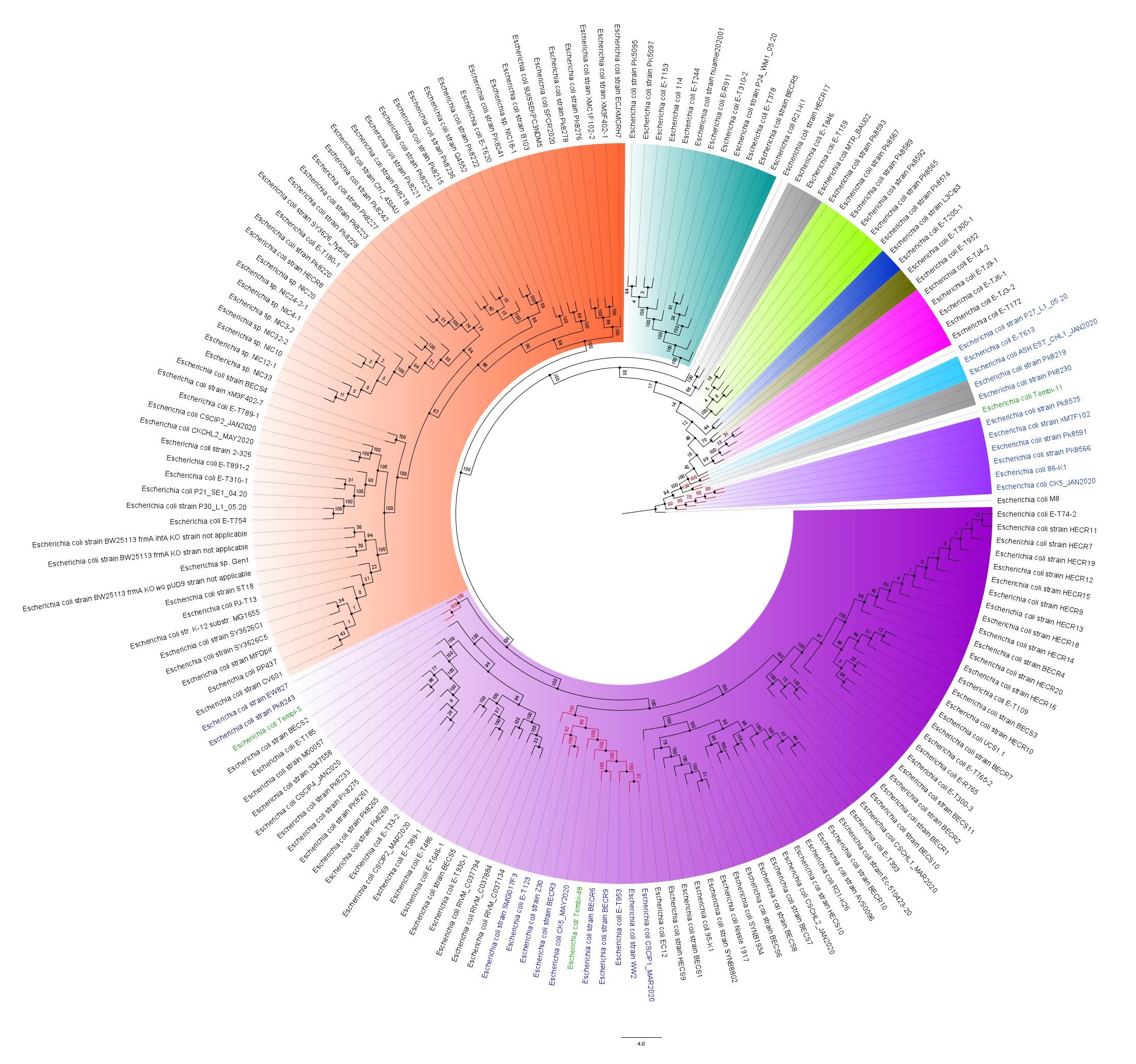

### Fig. S4

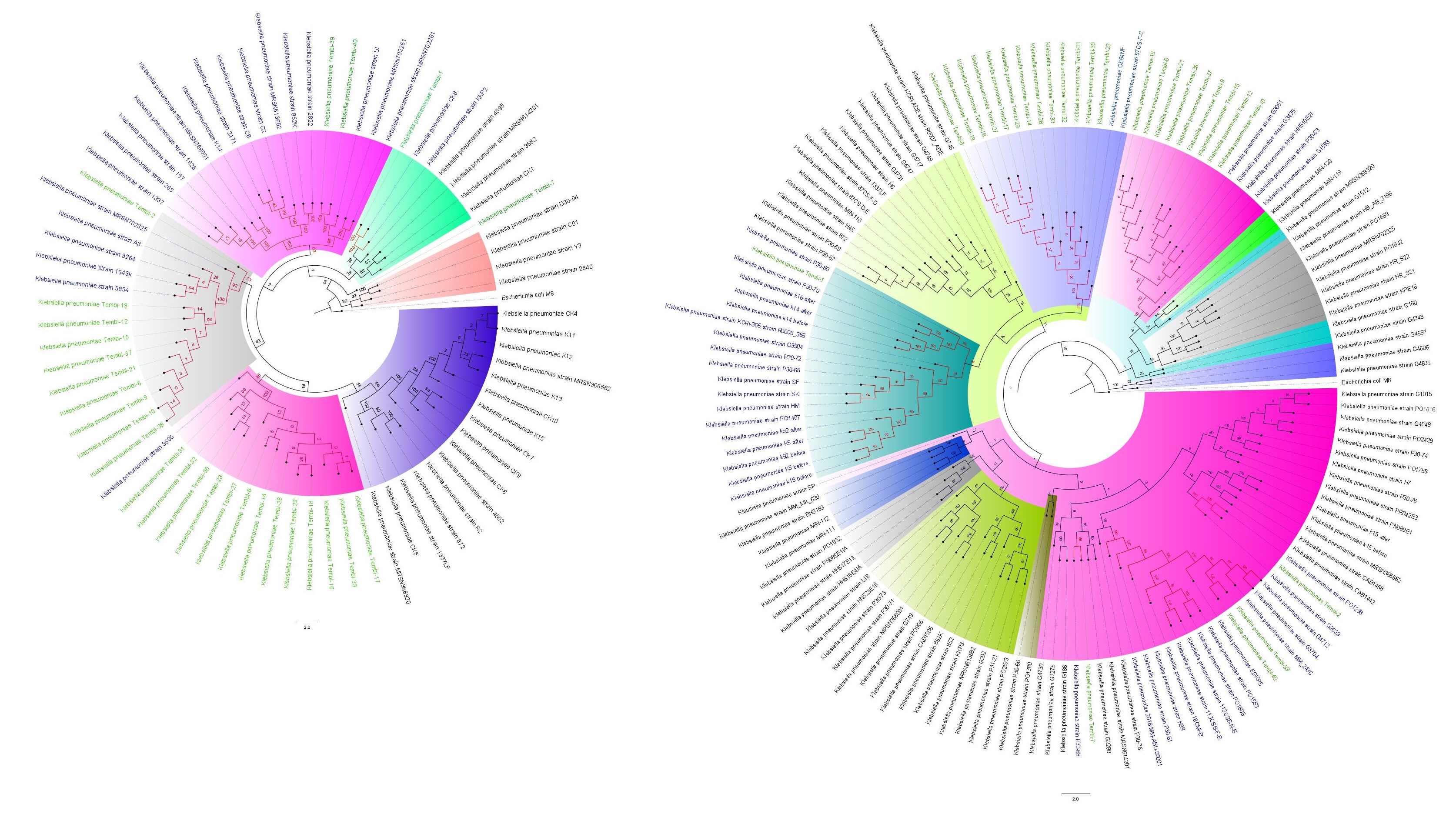

### Fig. S5A

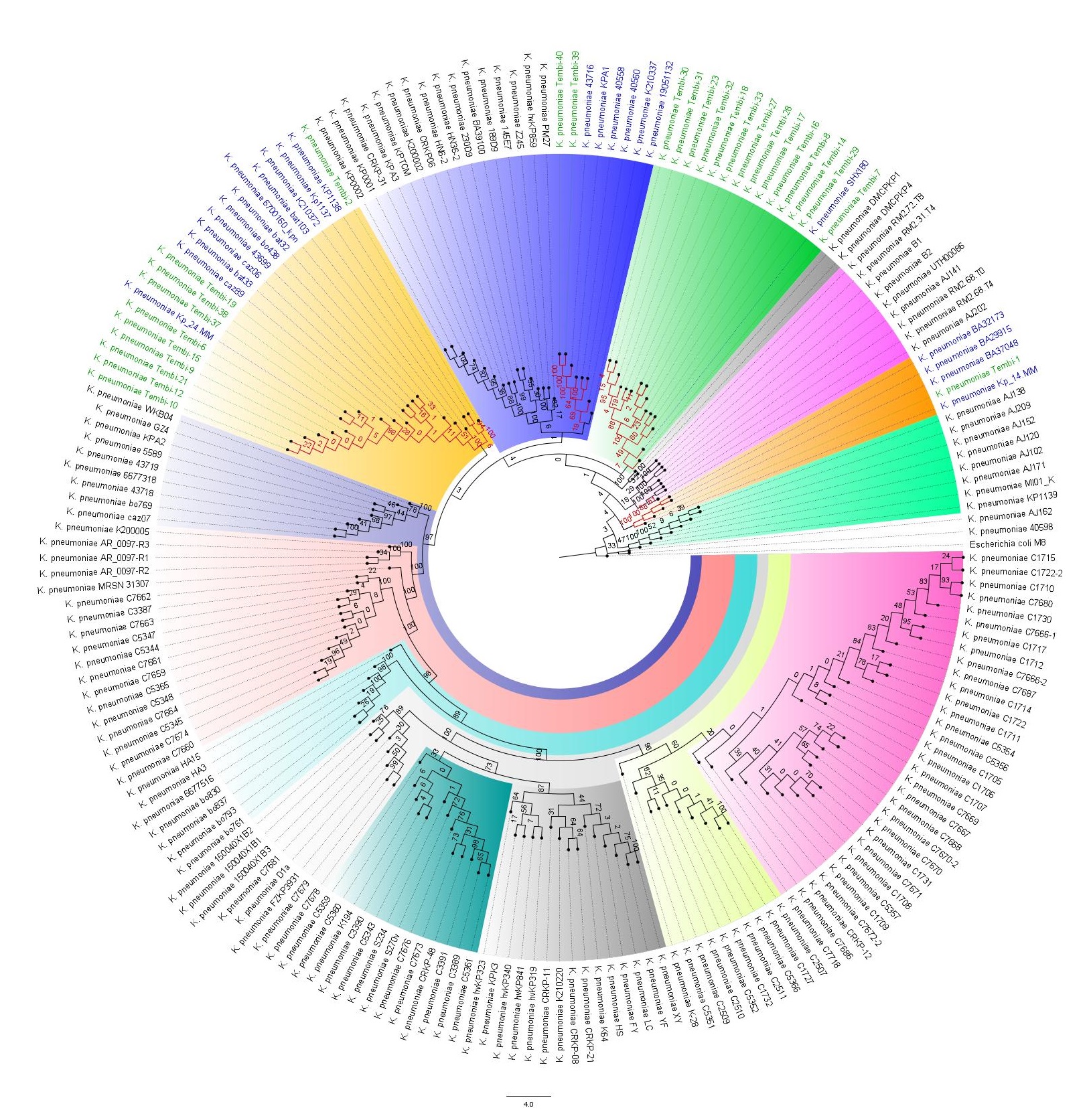

### Fig. S5B

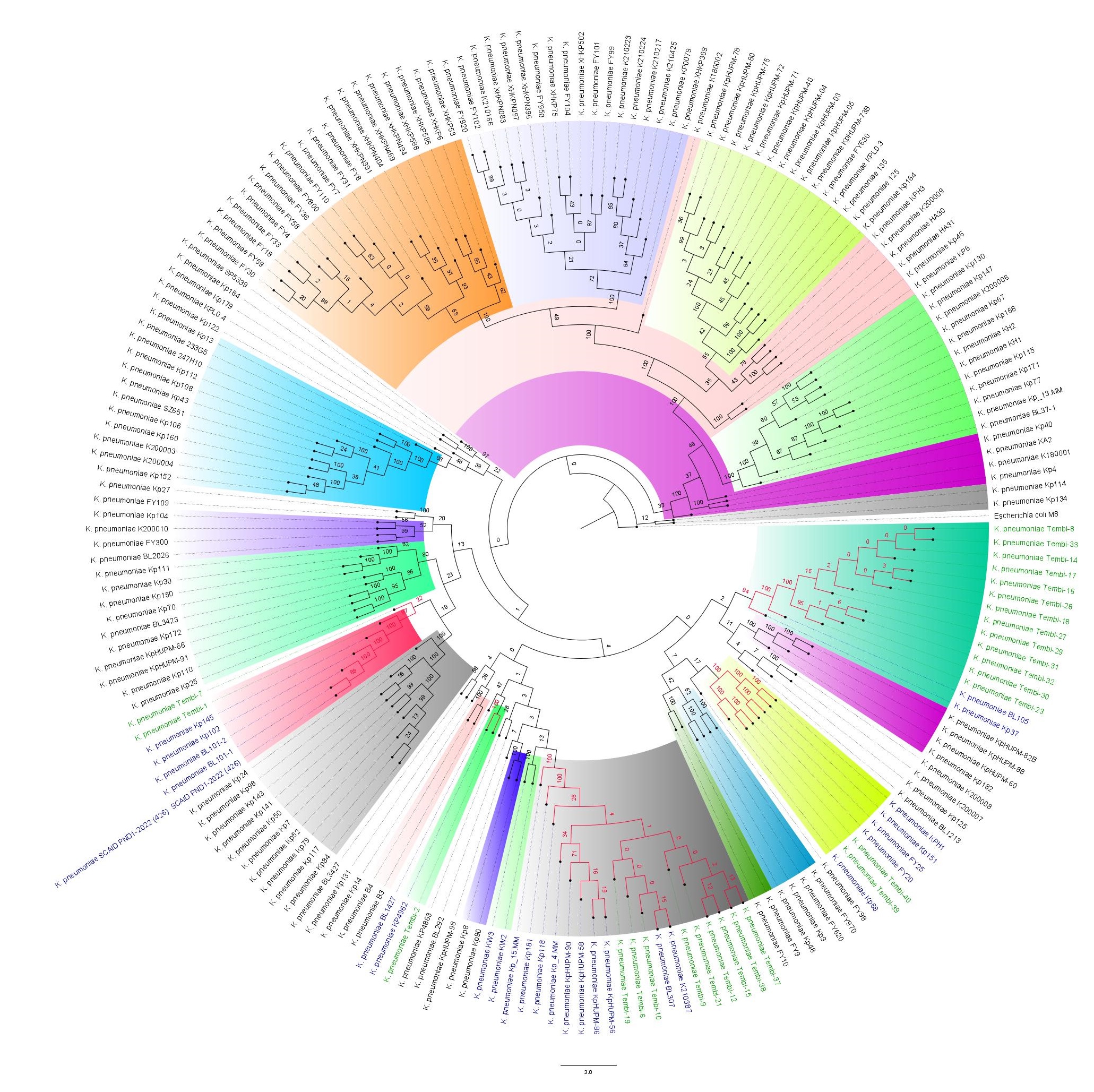

### S6

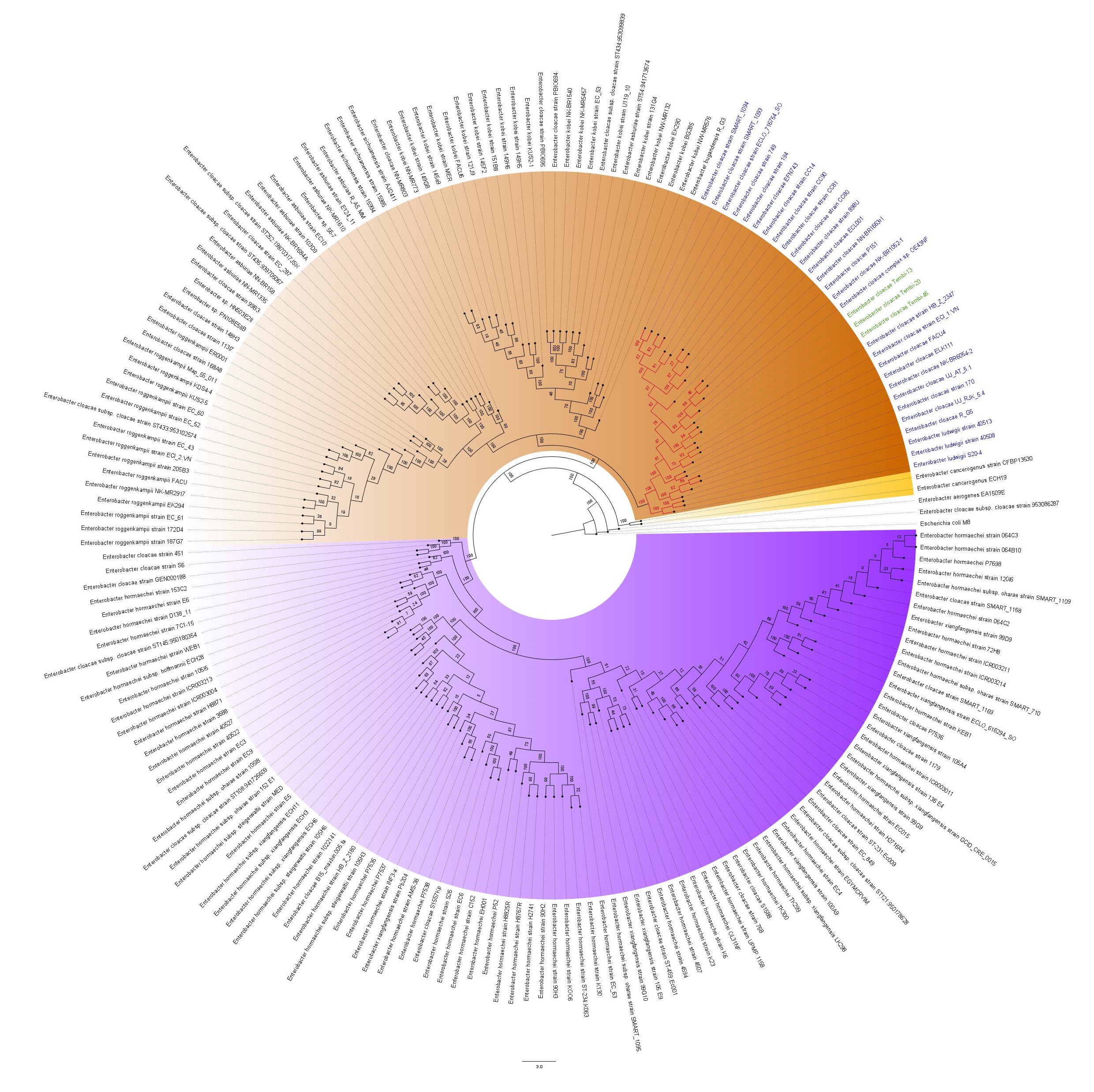

### S7

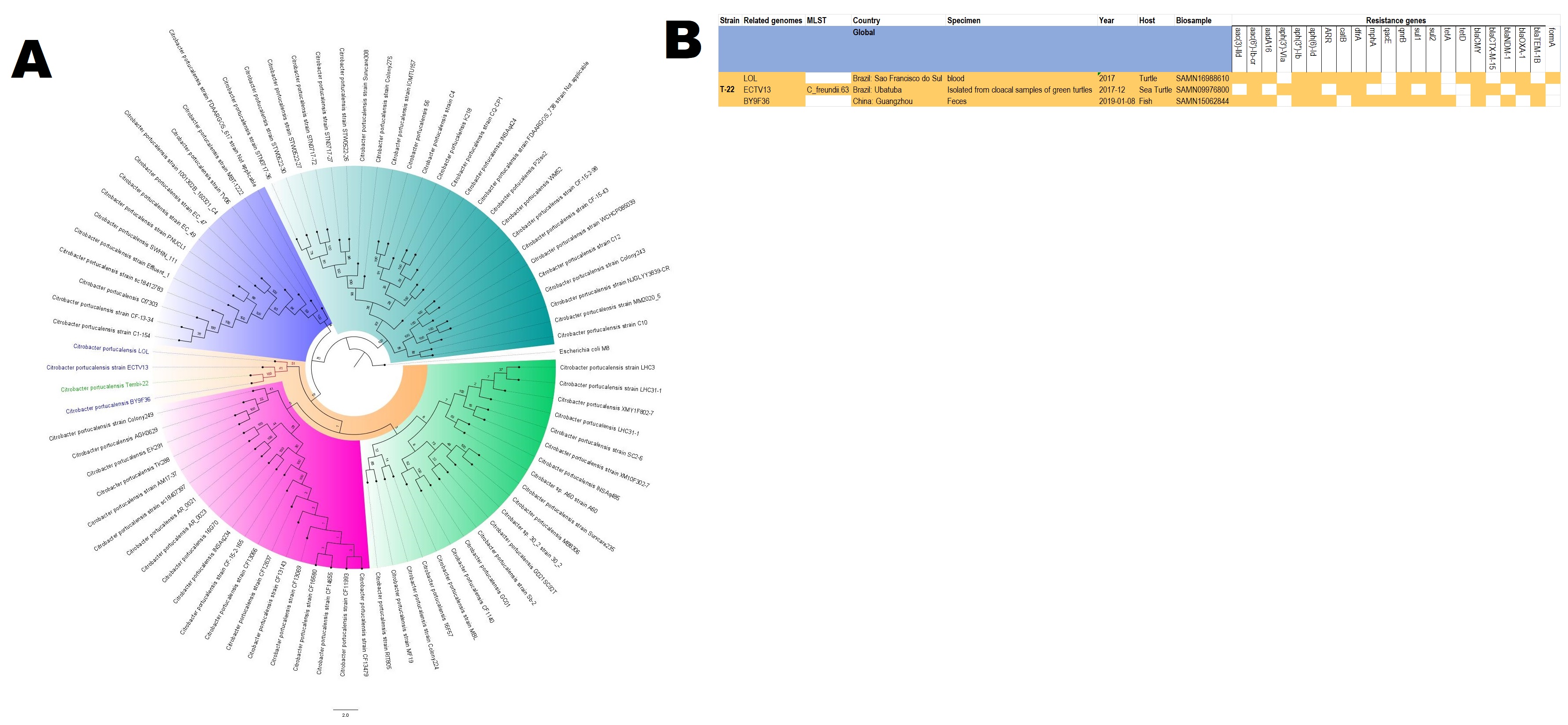
